# Behavioral clusters revealed by end-to-end decoding from microendoscopic imaging

**DOI:** 10.1101/2021.04.15.440055

**Authors:** Chia-Jung Chang, Wei Guo, Jie Zhang, Jon Newman, Shao-Hua Sun, Matt Wilson

## Abstract

*In vivo* calcium imaging using head-mounted miniature microscopes enables tracking activity from neural populations over weeks in freely behaving animals. Previous studies focus on inferring behavior from a population of neurons, yet it is challenging to extract neuronal signals given out-of-focus fluorescence in endoscopic data. Existing analysis pipelines include regions of interest (ROIs) identification, which might lose relevant information from false negatives or introduce unintended bias from false positives. Moreover, these methods often require prior knowledge for parameter tuning and are time-consuming for implementation. Here, we develop an end-to-end decoder to predict the behavioral variables directly from the raw microendoscopic images. Our framework requires little user input and outperforms existing decoders that need ROI extraction. We show that neuropil/background residuals carry additional behaviorally relevant information. Video analysis further reveals an optimal decoding window and dynamics between residuals and cells. Critically, saliency maps reveal the emergence of video-decomposition across our decoder, and identify distinct clusters representing different behavioral aspects. Together, we present a framework that is efficient for decoding behavior from microendoscopic imaging, and may help discover functional clustering for a variety of imaging studies.

## 1 Introduction

Tracking the activity of large neuronal populations in awake and behaving animals is crucial to understand the neural computation underlying cognitive functions. Recent advances in *in vivo* calcium imaging technology have nurtured studies of the neural circuits underlying sensory perception (Glas et al., 2019; Mittmann et al., 2011; Rothschild et al., 2010; Wang et al., 2020; Yoshida and Ohki, 2020), motor control (Ebina et al., 2018; Huber et al., 2012; Klaus et al., 2017), innate behaviors (Betley et al., 2015; Evans et al., 2018; Jennings et al., 2015), and a wide variety of cognitive behaviors such as decision making (Harvey et al., 2012; Pinto and Dan, 2015; Tanimoto et al., 2017), as well as learning and memory (Grewe et al., 2017; Roberts et al., 2017; Yu et al.,2017; Ziv et al., 2013). Specifically, a miniaturized head-mounted microscopy combined with a microendoscopic lens enables deep brain imaging of freely moving animals (Cai et al., 2016; Flusberg et al., 2008; Ghosh et al., 2011). Although two-photon miniature microscopes (Helmchen et al., 2001; Sawinski et al., 2009; Zong et al., 2017) have been developed to achieve optical sectioning, they have not been as widely adopted due to technical limitations that include slower acquisition speed, significant motion artifacts, and optical limitations of the fibers on delivering 920nm femtosecond laser pulses. On the other hand, microendoscopic imaging using a single-photon light source suffers from large background fluctuations due to out-of-focus fluorescence. Toward achieving the goal of decoding sensory stimuli or behavioral variables from the recordings, how to properly extract cellular signals from these low-contrast images has been a key step in analysis pipelines for calcium imaging (Lu et al., 2018; Pnevmatikakis, 2019; Zhou et al., 2018).

A typical analysis pipeline (Figure 1a) includes the following steps: First, registering image frames by correcting motion artifacts. Oftentimes, brain motion within the field of view (FOV) can be non-rigid along all directions and lead to frame distortions. To correct these motion artifacts, either a rigid template matching method (Dubbs et al., 2016; Kaifosh et al., 2014) or a non-rigid registration method (Lu et al., 2018; Pnevmatikakis and Giovannucci, 2017) is applied. Second, extracting neuronal signals through estimating spatial filters and their corresponding temporal traces. Common methods can be divided into three classes: semi-manual region of interest (ROI) analysis (Klaus et al., 2017; Pinto and Dan, 2015), principal component analysis/independent component analysis (PCA/ICA) (Mukamel et al., 2009; Reidl et al., 2007), and constrained nonnegative matrix factorization (CNMFe) (Pnevmatikakis et al., 2016; Zhou et al., 2018). Next, inferring spike times from extracted fluorescence traces. To simplify spike inference, fluorescence traces can be viewed as spike trains convolved with a kernel with asymmetrical rising and falling kinematics. Various algorithms (Jewell and Witten, 2018; Pnevmatikakis et al., 2016; Vogelstein et al., 2010; Yaksi and Friedrich, 2006; Zhou et al., 2018) have been proposed to solve this deconvolution problem.

**Figure 1:**
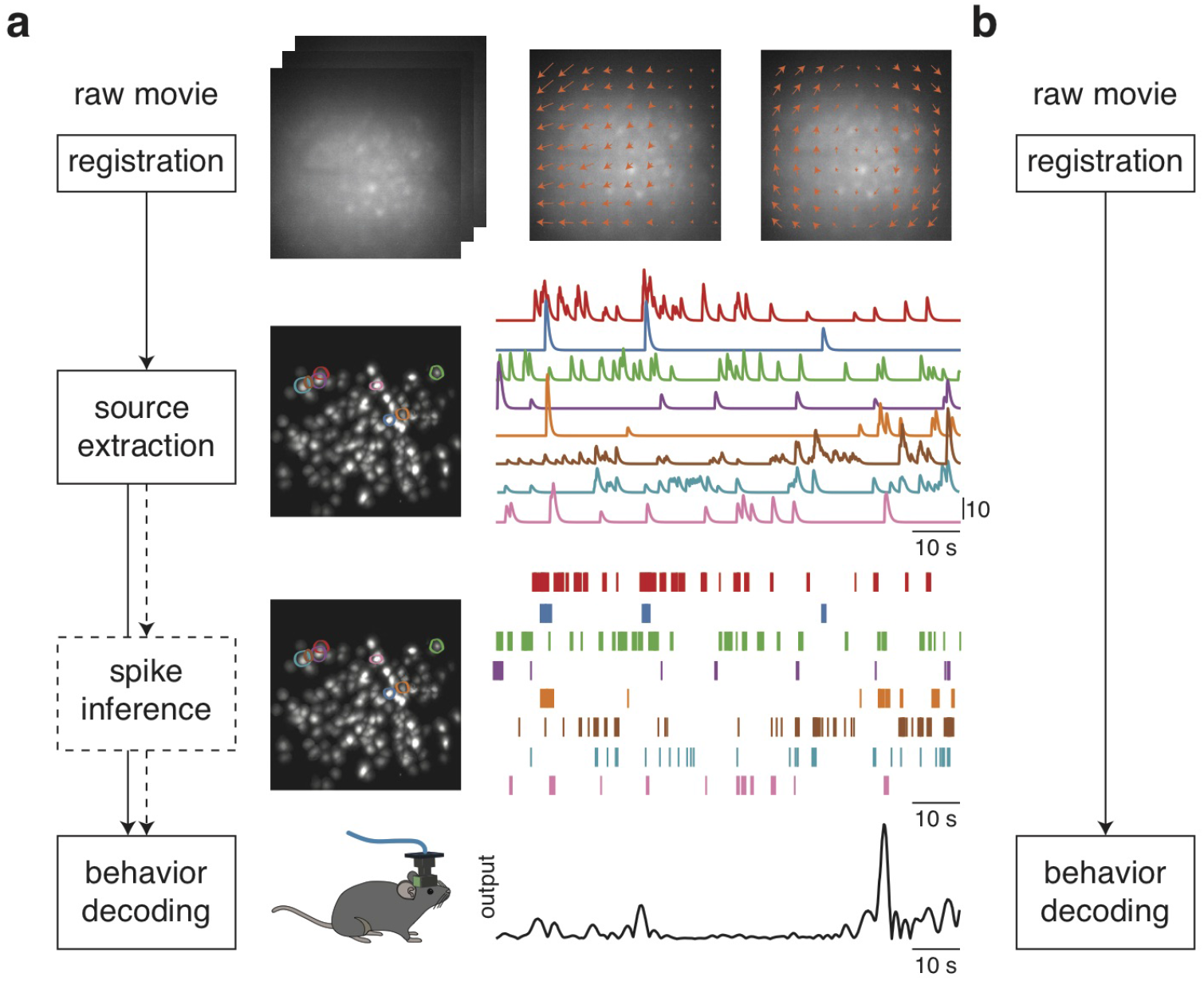
Typical and proposed analysis pipelines for one-photon imaging data. (**a**) Typical analysis pipeline. The data is first processed to remove motion artifacts by estimating a motion field from each data frame to a template (*top*). Due to limited axial resolution and low contrast in microendoscopic imaging, neurons can appear spatially overlapped and often require a video decomposition algorithm such as CNMFe (see Materials and Methods). After removing background signals, the locations of the neurons in the field of view (FOV) and their activities are extracted (*middle*). Spiking activity can then be inferred from each fluorescence trace. Finally, either spikes or fluroscence traces were used to decode the animal’s behavior (*bottom*). Different steps of the pipeline are displayed on one animal in our experiment. (**b**) Proposed pipeline. Animal behavior is directly decoded from raw movies after removing motion artifacts.

Finally, decoding behavioral variables or sensory stimuli from either extracted fluorescence traces or inferred spike trains. Common decoders including population vector reconstruction (Georgopoulos et al., 1986; Salinas and Abbott, 1994), Wiener filter (Carmena et al., 2003), template matching (Wilson and McNaughton, 1993; Zhang et al., 1998)), Bayesian paradigm (Brown et al., 1998; Sanger, 1996; Wu et al., 2006; Zhang et al., 1998), and machine-learning models (e.g., support vector machine (SVM), k-nearest neighbors (kNN), Fisher’s linear discriminant analysis (LDA), multilayer perceptron (MLP), recurrent neural network (RNN)) have been implemented in various studies (Glaser et al., 2020; Pereira et al., 2009; Quiroga and Panzeri, 2009).

Here we propose an end-to-end decoding paradigm to directly extract behavioral information from raw single-photon microendoscopic data (Figure 1b), and assess using the problem of predicting kinematics from calcium imaging of hippocampal CA1 in freely foraging mice (Figure 2a, b). We hypothesize that the neuropil/background, which was discarded in most analysis pipelines, may encode the animal’s position and/or speed. Instead of making assumptions on the format of relevant signals, our paradigm leverages recent progress in deep learning to determine which components of imaging data are most informative without a need of sophisticated preprocessing. Our decoder outperforms decoding methods which use extracted neuronal signals. The results compare encoding capacity between putative neurons and the background residuals, and identify contribution across timepoints. By examining the saliency maps, we find video decomposition naturally emerges in our decoder. Our findings suggest neuropil/background as additional information encoder in microendoscopic recordings, and provide a new way to identify neural ensembles representing distinct aspects of spatial navigation.

**Figure 2:**
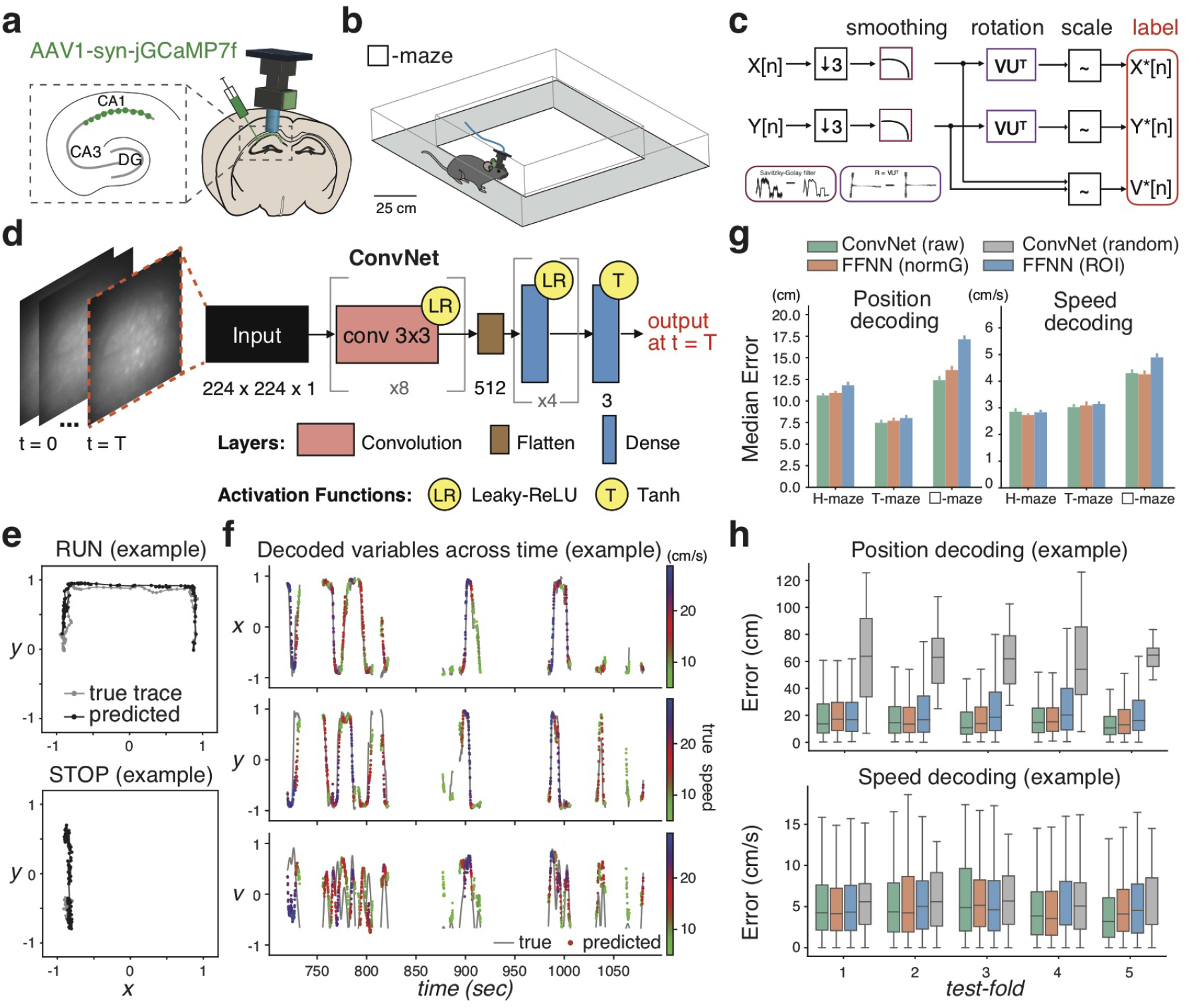
Accurate decoding of positions and running speeds from raw microendoscopic images. (**a-d**) Schematic of data preparation and model architecture. Details see Materials and Methods. (**a**) Microendoscopic imaging in the CA1 expressing jGCaMP7f. (**b**) An example of H-maze exploration experiment. Each maze has a length of about 1 m. (**c**) Animal positions (*x, y*) were aligned to imaging data, downsampled to 10 Hz, smoothed and then rotated to reduce their correlations. Finally, positions and speeds (*x, y, v*) were scaled to [-1, 1]. (**d**) ConvNet architecture. At each timepoint, an image was fed to a deep network consisting of 2D convolutional layers with leaky-ReLU activations, followed by dense layers with a regression head to decode position (*x, y*) and instantaneous speed (*v*) of the animal. (**e**) Example 10-sec trajectories of true (gray) and decoded (black) positions from an animal in a □-maze exploration experiment. Top: A 10-sec trajectory when the animal was running. Bottom: A 10-sec trajectory when the animal was stationary or running at a speed below 5 cm/s. (**f**) Decoded variables across time in an example test set. Gray line denotes true traces. Dots denote predicted traces, colored based on running speeds. Average error in decoding x, y, and v: 7.84 cm, 12.70 cm, 6.23 cm/s, respectively. (**g**) Model performance across all datasets. Left: Our ConvNet model had less median error in predicting positions (10.17 ± 0.43 cm) than baseline decoders. They were all better the chance level (48.57 ± 2.67 cm; *p* < 1*e* − 5). Right: Speed decoding in our ConvNet model (3.39 ± 0.14 cm/s) was comparable to a decoder trained on neuronal signals (3.36 ± 0.14 cm/s; *p* = 0.78), and better than a decoder trained on ROI signals (3.62 ± 0.18 cm/s; *p* < 0.05). All were better than the chance level (4.66 ± 0.19 cm/s; *p* < 1*e* − 5). (**h**) Error distribution of position (*top*) and speed (*bottom*) decoding in an example dataset across test-folds. All p-values were obtained by Wilcoxon signed-rank test.

## 2 Previous Work

In a typical analysis pipeline, how neuronal signals are extracted and structured is a key determinant with single-photon microendoscopic data. Unfortunately, existing source extraction methods have several limitations. Take semi-manual ROI analysis (Klaus et al., 2017; Pinto and Dan, 2015) as an example, neuropil contamination from out-of-focus and scattered fluorescence is often inadequately removed and require explicit assumptions on neuropil structure. Plus, signals of spatially overlapped neurons cannot be demixed. In addition, manual ROI annotation can be highly variable with inter-labeler disagreement and laborious for large-scale recordings, whereas thresholding an activity map, which indicates changes of the fluorescence value across time at each pixel, often leads to a high miss rate for ROI detection. As for PCA/ICA analysis (Mukamel et al., 2009; Reidl et al., 2007), it ignores the fact that neurons usually have signal correlations and noise correlations (Averbeck et al., 2006). Moreover, it is a linear method and cannot properly demix signals of spatially overlapped neurons. Recently, CNMFe (Zhou et al., 2018) has gained popularity given its superior ability in demixing and denoising neuronal signals from microendoscopic data. However, it depends on sophisticated hyperparameter tuning which requires prior knowledge on the number of neurons and their sizes. Additionally, it is sensitive to parameter initializations and has non-unique solutions for ROI identification. Specifically, all these methods have detection errors, which lead to either introducing extra bias or throwing away relevant information. On top of issues above, these algorithms are often time-consuming.

A recent study developed a new signal extraction pipeline for single-photon microendoscopic imaging (MIN1PIPE) (Lu et al., 2018) to improve ROI identification. By incorporating a two-component Gaussian mixture model (GMM) and a trained recurrent neural network (RNN), fluctuations unlikely to be calcium transients are removed, leading to a cleansed set of ROI seeds. This method reduces the need to set many hyperparameters heuristically. However, it is computationally expensive and requires extra manual judgement for training to classify calcium transients. Recognizing the challenge of obtaining highly preprocessed data for decoding, another study developed Deepinsight (Frey et al., 2019), a deep-learning framework, to infer behaviors from wide-band neural data (e.g., raw electrophysiology recordings without spike sorting). However, when it comes to calcium imaging data, this study still requires extracting temporal traces from identified ROIs, before converting them into frequency representations which are provided to the decoder as input.

An end-to-end decoder is widely used in many machine-learning domains, but has never been applied in the neuroscience field to our knowledge. Here we aim to directly extract behavioral information from raw microendoscopic recordings, and examine its learned representations.

## 3 Results

To evaluate how well behavioral information can be directly extracted from microendoscopic imaging data by our proposed decoding paradigm (Figure 1b), we expressed Ca^2+^ indicator jGCaMP7f in the dorsal hippocampus of mice, implanted a GRIN lens right above the CA1 region (Figure 2a), and recorded calcium activity in the CA1 through a miniature head-mounted single-photon microscope while the animal was freely navigating different mazes (Figure 2b and Supplementary Figure 1a). Imaging data and the animal’s behavioral information were simultaneously recorded, synchronized at a sampling rate at 30 Hz.

To generate required inputs and labels for the decoder model, we first preprocessed behavioral data in the following steps. Both images and animal positions (*x,y*) were downsampled to 10 Hz to reduce memory consumption. A Savitzky-Golay filter was applied to smooth position coordinates and obtain running speeds (*v*). Afterwards, position coordinates were rotated to encourage orthogonalization between output labels. Finally, all the variables (*x, y, v*) were scaled to [-1, 1] (Figure 2c). As for image inputs, the field of view (FOV) was cropped into 224 by 224 pixels, with values being scaled as well (see Materials and Methods).

Prior to model training, we split each dataset into a training set and a test set. By segmenting each session into five sequential periods, we selected one of them as the test set. In the literature, a rolling forecasting strategy for train/test split is often applied to time-series data, by assuming that distributions are time-invariant, which might not be appropriate if the animals change their behavioral signatures over time.

### 3.1 Accurate decoding of positions and running speeds from raw microendoscopic data

To directly decode positions and running speeds of the animal from raw imaging data, we built a convolutional regression model (ConvNet) (Figure 2d). At each timepoint, a single frame was fed to a network consisting of 2D convolutional layers with leaky-ReLU activations, followed by dense layers. A Tanh operation was applied to the last layer. The hyperparameters of the model were searched and determined based on the validation set. Results showed that our ConvNet model accurately decoded the positions of the animals as well as their running speeds from unprocessed microendoscopic images, across different experiments (Figure 2e-h and Supplementary Figure 2a-c). By comparing decoding performance with a feedforward neural network (FFNN) trained on CNMFe-denoised neuronal signals, i.e., FFNN(normG), and a model trained on background-removed ROI signals, i.e., FFNN(ROI) (see Materials and Methods), we found that in terms of predicting positions (Figure 2g, left), our ConvNet model had less median decoding error (10.17 ± 0.43 cm) than the FFNN(ROI) model (12.32 ± 0.72 cm; Wilcoxon signed-rank test: stats = 54, *p* < 0.001) and the FFNN(normG) model (10.74 ± 0.49 cm; stats = 135, *p* < 0.05). These models were all significantly better than the chance level (48.57 ± 2.67 cm; *p* < 1*e* − 5). As for predicting running speeds (Figure 2g, right), the median decoding error of our ConvNet model (3.39 ± 0.14 cm/s) was less than the FFNN(ROI) model (3.62 ± 0.18 cm/s; stats = 121, *p* < 0.05), and comparable to the FFNN(normG) model (3.36 ± 0.14 cm/s; stats = 219, *p* = 0.78). These models were also better than the chance level (4.66 ± 0.19 cm/s; *p* < 1*e* − 5).

### 3.2 Background residuals encode behavioral information

Given that our ConvNet model trained on raw microendoscopic recordings was able to decode positions and speeds better than a decoder trained on CNMFe-extracted neuronal signals, we hypothesized that neuropil/background residuals, often discarded in imaging studies, encode additional behavioral information than cell somata.

To evaluate how much behavioral information is embedded in different components in microendoscopic imaging, we generated several image sets (Figure 3a), including Clean images composed with cell somata only, Residual images contained signals after subtracting neuronal signals from raw data, and Hollow A images where spatial footprints of detected cells were occluded (see Materials and Methods). As revealed by an example test set (Figure 3b), both models trained on Raw and Clean images were able to decode behavioral variables (*x, y, v*) across time. Surprisingly, models trained on Residual images were still able to predict behavioral traces (Figure 3b, c). This result was not unique to a single test set. We evaluated all the datasets from different maze exploration experiments (Figure 3d), and found decoding positions from Residual images had a median error (14.93 ± 1.04 cm) comparable to decoding from Clean images (11.03 ± 0.57 cm; stats = 165, *p* = 0.16). Models trained on Hollow ROI (13.46 ± 0.64 cm) and Hollow A (15.13 ± 0.69 cm) images had significantly less median decoding error than the chance level (48.57 ± 2.67 cm; *p* < 1*e* − 5).

**Figure 3:**
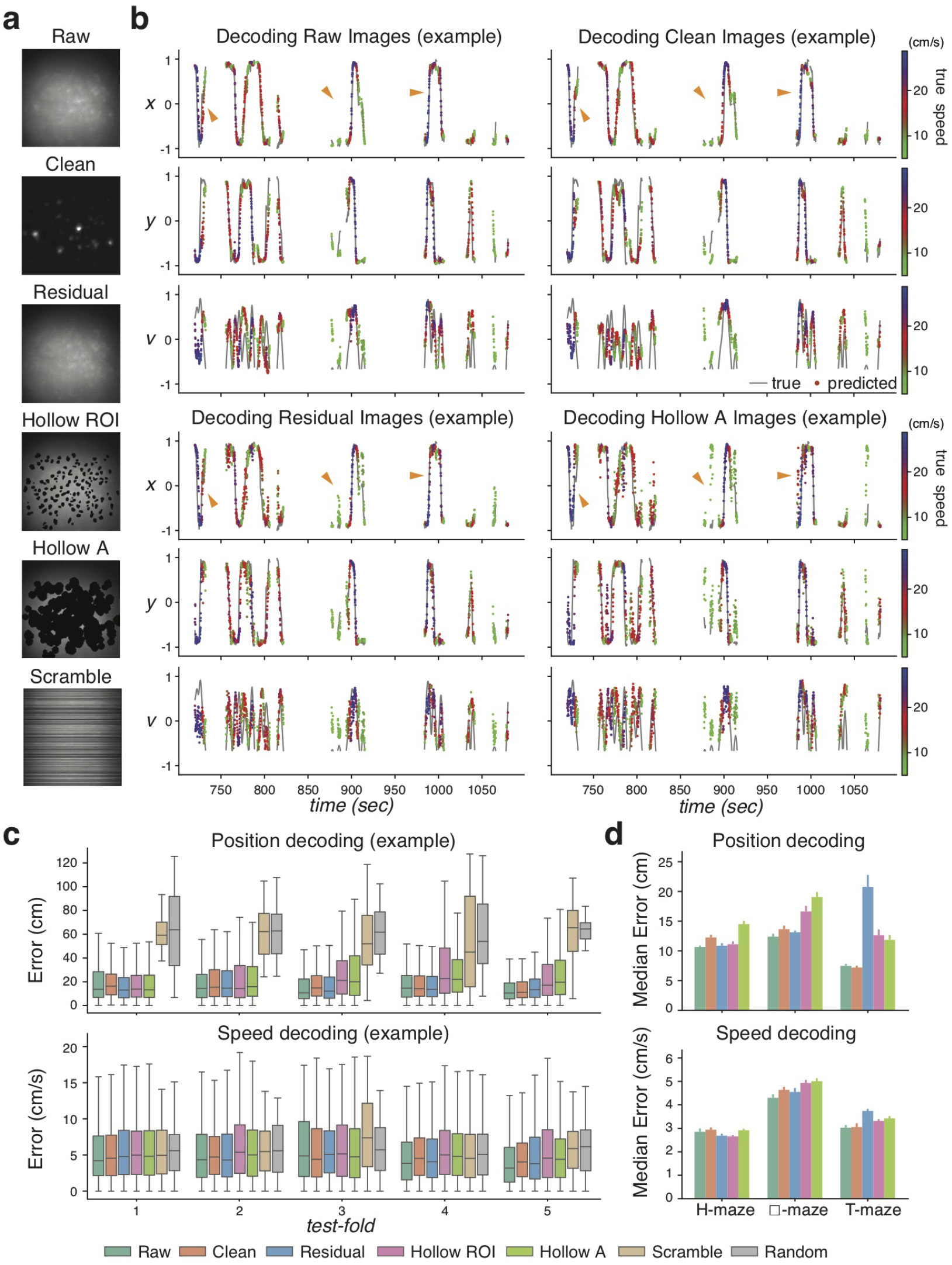
Neuropil/background residuals encode behavioral information. (**a**) Data generation for evaluating information embedded in different components in one-photon calcium imaging. Clean: Images composed with denoised neuronal signals only. Residual: Residuals from subtracting Clean from Raw images. Hollow ROI: Residuals with ROI removed. Hollow A: Residuals with CNMFe detected cell footprints removed. Details see Materials and Methods. (**b**) Examples of variables decoded by using images prepared in **a** from the same dataset. Orange triangles marked timepoints where we can easily see prediction differences between models. Gray line denotes true traces. Dots denote predicted traces, colored based on running speeds. (**c**) Error distribution of position (top) and speed (*bottom*) decoding in an example dataset (same as **b**) across test-folds. (**d**) Model performance across all datasets. Top: Decoding positions using Residual images (14.93 ± 1.04 cm, blue) had more median error than using Raw images (10.17 ± 0.43 cm, green; *p* < 0.01), but comparable to using Clean images (11.03 ± 0.57 cm, orange; *p* = 0.16). Both Hollow ROI (13.46 ± 0.64 cm, pink) and Hollow A (15.13 ± 0.69 cm, lime) images had significantly less median decoding error than the chance level (48.57 ± 2.67 cm; *p* < 1*e* − 5). Bottom: Decoding speeds using Residual images (3.66 ± 0.15 cm/s, blue) had more median error than using Raw images (3.39 ± 0.14 cm/s, green; *p* < 0.05), but similar to using Clean images on average (3.54 ± 0.16 cm/s, orange; *p* = 0.46). Both Hollow ROI (3.63 ± 0.18 cm/s, pink) and Hollow A (3.78 ± 0.17 cm/s, lime) images had significantly less median decoding error than chance level (4.66 ± 0.18 cm/s; *p* < 1*e* − 5). All p-values were obtained by Wilcoxon signed-rank test.

If there was no extra information embedded in residuals compared to identified neurons, we would have observed a similar decoding performance between models trained on Raw and Clean images. Instead, decoding positions from Raw images, with no information excluded, had less median error (14.93 ± 1.04 cm) than decoding from Clean images (11.03 ± 0.57 cm; stats = 102, *p* < 0.01). In some cases, a model trained on Residual images was able to decode positions even better than using Clean images (e.g., across H-maze exploration datasets: Residual: 10.87 ± 0.42 cm, Clean: 12.25 ± 0.43 cm; stats = 4, *p* < 0.05). These results suggest that additional position information can be embedded in neuropil/background residuals.

### 3.3 Incorporating multiple frames into the model input improves decoding

Previous results were decoded and examined at a frame-by-frame level. Given the relatively slow calcium dynamics, we further hypothesize that incorporating multiple frames into the input improves decoding performance. To test this hypothesis, we modified the ConvNet architecture to incorporate multiple frames in the input (see Materials and Methods for details).

When there was no constraint on convolutional kernels, inputs with 51 frames corrected temporal mismatch in predictions compared to single-frame input (Supplementary Figure 3a). Across all datasets, position and speed decoding improved significantly by incorporating multiple frames into the input (Supplementary Figure 3b). Specifically, using 11 frames (1-sec time window) had least median error in position prediction (single-frame: 10.17 ± 0.43 cm; 1-sec time window: 7.50 ± 0.60 cm; stats = 5, *p* < 1*e* − 5).

We further modified the architecture by adding a constraint on convolutions across the temporal dimension (Figure 4a). Specifically, each frame went through the same block of convolutional layers whose weights were shared across frames. By learning attention weights attributed to these frames, information about the behavioral state was extracted from different timepoints. A similar decoding improvement by incorporating multiple frames into the input was observed (Figure 4b). By examining the distribution of learned attention weights for an input with 51 frames, we found that the imaging frame contributed most to decoding the behavioral state was delayed (Figure 4c), suggesting future imaging frames carry more information about the behavioral states of the animal.

**Figure 4:**
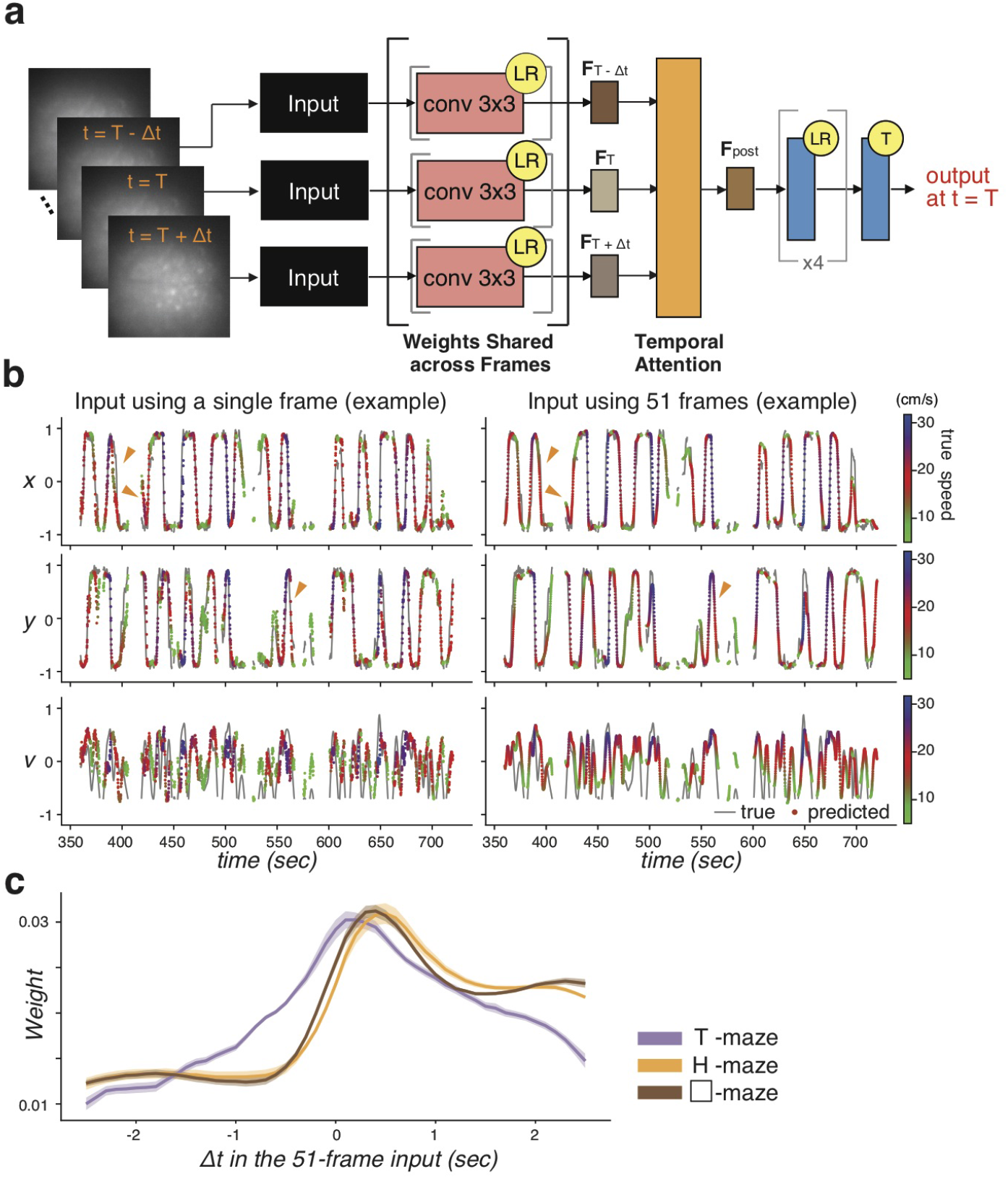
Incorporating multiple frames into the model input improves decoding. (**a**) Schematic diagram of ConvNet modified to incorporate multiple frames into the input. Each frame in the input goes through a block of convolutional layers whose weights are shared across frames, and generates a frame-specific feature (e.g., *F*_*T*-Δ*t*_, *F_T_*, and *F*_*T*+Δ*t*_). The attention layer learns weights attributed to frames at different timepoints in the input. The weighted-average feature (*F_post_*) is fed into the dense layers. (**b**) Examples of variables decoded from the same dataset, using a single-frame input (*left*) and 51 frames (*right*) when convolutions were shared across frames. Orange triangles mark timepoints where we can easily see prediction differences between models. The gray line denotes true traces. Dots denote predicted traces, colored based on running speeds. (**c**) The distribution of learned attention weights for an input with 51 frames (Δ*t* = ± 2.5 seconds).

### 3.4 Distinct functional ensembles are identified from raw microendoscopic images

One challenge in using a deep-learning model is to interpret what features are learned from the data. Our ConvNet model was able to decode the animal’s kinematics, but it was unclear how behavioral information was extracted from the microendoscopic images by the decoder.

To understand which information was extracted from the raw imaging data, we modified the Grad-CAM algorithm (Selvaraju et al., 2017) to identify saliency maps across different layers in the model (Figure 5a). Specifically, the saliency map of a target unit in a specific layer was estimated by a gradient-weighted combination of feature maps (see Materials and Methods for details). If the ConvNet decoder utilized the former learning strategy, the saliency maps would demonstrate uniform scattered patterns. On the other hand, if the latter strategy was used, the saliency maps were likely to identify ensembles composed with distinct combinations of cells and background features. We found that in the early stage of the training, global background fluorescence were already segmented from and neuronal signals across layers, and the edges of cell clusters were partially detected in the middle convolutional layers (Figure 5b, top). When the model was well-trained (i.e., after 26000 steps), auto-decomposition was emerged across different layers (Figure 5b, bottom). Specifically, global background signals were segmented out in early layers of the model, but distinct ensembles of cells and background features were identified in the later layers of the model.

**Figure 5:**
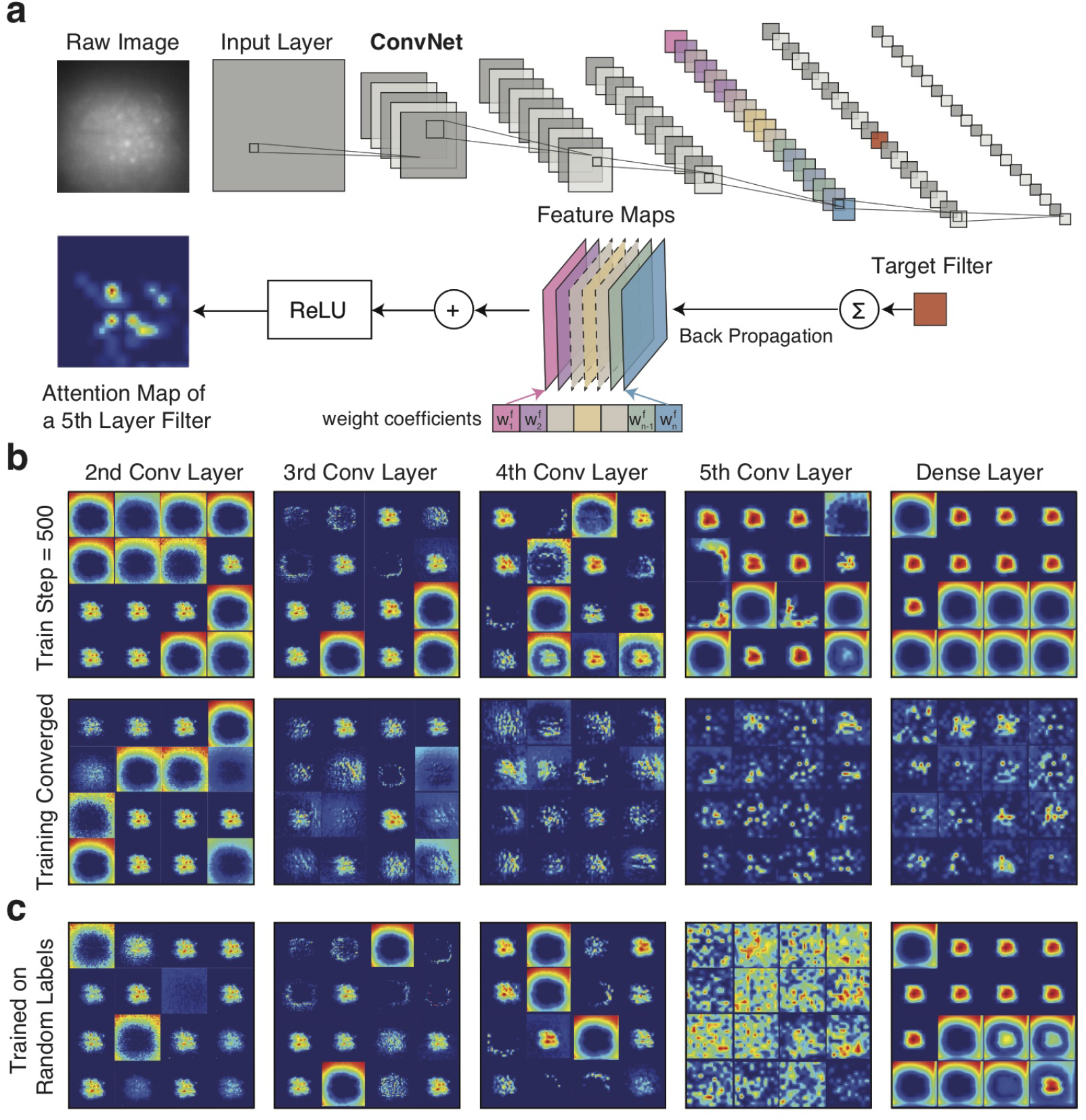
Emergence of auto-decomposition and functional clusters. (**a**) Schematic diagram of adapted Grad-CAM algorithm. We first back-propagate gradients of a target filter at a specific layer to the closest previous convolutional layer. The gradient-weighted combination of feature maps followed by a ReLU operation gives the estimated saliency map. Details see Materials and Methods. (**b**) saliency maps across different model layers. Each square box corresponds to saliency map of a typical unit/filter in that layer. For each layer, a example subset of saliency maps were shown. Top: When the model was trained with 500 steps only, global background fluorescence and signals from cell somata or local neuropils were already segmented across layers, and object edges were detected based on brightness in the middle convolutional layers. Bottom: When the model was well-trained, auto-decomposition was emerged across different layers. In early convolutional layers, similar global background signals were segmented out, whereas distinct ROIs were identified in the later convolutional layers. (**c**) saliency maps across different model layers when output labels were randomly assigned. A different decomposition process was formed.

One might argue that a convolutional model can naturally learn object identification by extracting morphological features in an image, so the emergent decomposition across model layers is simply a byproduct of a convolutional neural network. To test against this hypothesis, we compared saliency maps in a model where output labels were randomly assigned. If the hypothesis was true, we would expect similar saliency maps. Instead, a different decomposition process was formed when the relationship between input images and output labels were removed (Figure 5c), suggesting auto-decomposition that emerged from the ConvNet decoder was functionally relevant.

### 3.5 Differential encoding of the maze topology across layers

Following previous findings that distinct functional ensembles were identified by the decoder. We further examined how the maze topology was represented in these ensembles. Previous studies have shown that the intrinsic dynamics of a neural ensemble often occupy a low-dimensional manifold within the high-dimensional state space (Archer et al., 2014; Churchland et al., 2012; Cueva et al.,2020; Mante et al., 2013). Here, the neural ensemble consist of the activity states of artificial units in the decoder. To extract the representations from attended units in each layer of the trained model, we projected their activity into a 2-dimensional state-space using Isomap (see Materials and Methods). To further explore how the maze topology was learned across different layers in the ConvNet model, we mapped each neural state to the animal’s true positions in the maze (Figure 6a, b). When the model was well-trained (Figure 6a), the projected activity accurately captured the intersection and each arm of the maze in the middle layers of the model, despite the distances between these neural states being unable to capture the actual maze topology. However, in the later layers of the model, projected neural states started to reflect the actual maze topology. We also visualized the ensemble representations in an early training stage whose validation error was twice of the fully trained model (Figure 6b), and showed that these ensemble representations in early layers were determined early in the training, whereas the behavioral topology was shaped at later layers throughout training.

**Figure 6:**
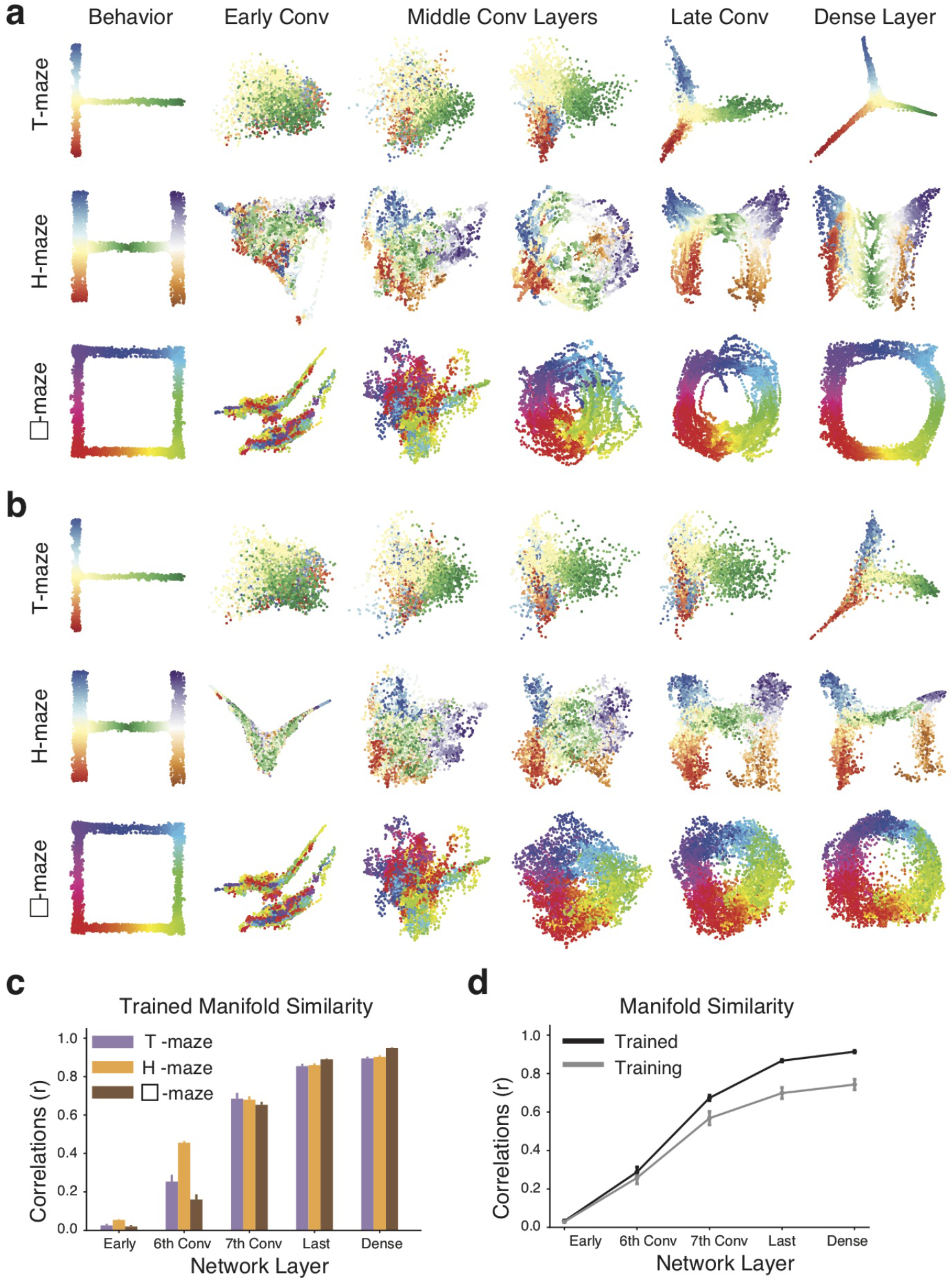
Encoding of behavioral topology in manifolds across model layers. (**a**) Ensemble representations from different layers of a trained model on a typical dataset in T-maze (*top*), H-maze (*middle*), and □-maze (*bottom*) experiments. We extracted the representations from attended units in each layer of the trained model, and projected their activities to a 2-dimensional state-space using Isomap. Each dot represents a behavioral state, colored by the animal’s true positions in the maze. The 1st Column: Position distributions from a typical animal. Other columns: The topology of the animal’s positions started to form at later layers. Details see Materials and Methods. (**b**) Ensemble representations from different layers of a model in an early training stage whose validation error was about twice. (**c**) Similarity between neural manifolds and behavioral topology. Similarity was defined as the correlation between pairwise distances in a neural manifold and pairwise distances in behavioral topology. Similarity increased significantly from the 6th convolutional layer to the 7th convolutional, and further increased in the last convolutional layer as well as the dense layer, across T-maze (purple), H-maze (gold) and □-maze (brown). (**d**) Comparison between neural manifolds in well-trained models (black) and models whose validation errors were about twice (gray). Across all datasets, manifold similarity was significantly higher in the trained models (black) than models under training (gray) at later layers (Wilcoxon signed-rank test: stats = 1, *p* < 1*e* − 5, whereas such difference was less yet still significant at earlier layers (Wilcoxon signed-rank test: stats = 98, *p* < 0.05).

To evaluate the similarity between each neural manifold and the maze topology, we computed the Pearson’s correlation between pairwise distances in the neural manifold and pairwise distances in the physical maze. As shown in Figure 6c and d, the similarity between each neural manifold and the maze increased from the early layers to the later layers of a trained model (Early: 0.03 ± 0.01; Conv6: 0.29 ±0.03; Conv7: 0.67 ± 0.01; Last: 0.87 ± 0.01; Dense: 0.91 ± 0.01), and was significantly higher compared to models in an early stage of the training (Early: 0.03 ±0.01, stats = 98, *p* < 0.05; Conv6: 0.26 ± 0.03, stats = 117, *p* = 0.05; Conv7: 0.57 ± 0.03, stats = 102, *p* < 0.05; Last: 0.70 ± 0.03; Dense: 0.74 ± 0.03, stats = 1, *p* < 1*e* − 5) across different maze exploration experiments.

Overall, these results reveal that the decoder represents the animal’s internal representations of the maze, by decomposing the maze topology across layers.

## 4 Discussion

### 4.1 Effect of data representations and model architectures

After showing that a ConvNet model can directly extract behavioral information from raw microendoscopic data, we evaluated how data representations affect decoding performance. Ideally, we would like output labels to approximate a uniform distribution for training a deep-learning model. However, the running speeds of the animals were heavily skewed toward lower values. We applied a log-transformation on speeds (Supplementary Figure 4a), and re-trained a subset of models. Results showed that the original ConvNet trained with linear speeds (11.49 ± 0.13 cm) had significantly less median error in predicting positions than the same architecture trained with log speeds (12.45 ± 0.14 cm; *p* < 1*e* − 5) (Supplementary Figure 4b, left). Similarly, the median error in predicting running speeds was significantly lower in the original model trained with linear speeds (3.76 ± 0.03 cm/s) than the model trained with log speeds (3.95 ± 0.04 cm/s; *p* < 1*e* − 5) (Supplementary Figure 4b, right). This result matched with studies suggesting neurons in the hippocampus encoded running speeds in a linear manner (Góis and Tort, 2018; Kropff et al., 2015), different from log-scale representations in the sensory cortex (Nover et al., 2005).

We further evaluated how different model architectures could affect the decoding performance with our datasets. The original ConvNet model predicted positions and speeds simultaneously (Supplementary Figure 5a, top). We hypothesized that speed decoding benefited from a joint training paradigm, and a separate decoding strategy could remove such benefits. We modified the model to have two separate decoders, one for position and the other for speed (Supplementary Figure 5a, bottom), and compared their performance in predicting positions and running speeds of the animal (Supplementary Figure 5b). Results showed that a model with two separate decoders improved position prediction (9.85 ± 0.42 cm) compared to the original model with simultaneous decoding (10.17 ± 0.43 cm; stats = 133, *p* < 0.05), but came at a cost in speed decoding, i.e., its median error (3.53 ± 0.16 cm/s) was more than the original model (3.39 ± 0.14 cm/s; stats = 131, *p* < 0.05).

We also did transfer learning with a ResNet-50 model and a MobileNet model (Supplementary Figure 6a), whose feature extraction parameters were pre-trained by the ImageNet and frozen during model training, whereas the parameters in the dense layers were updated throughout training (see Materials and Methods). We also built a feedforward neural network to extract features from raw microendoscopic images after a max pooling (Supplementary Figure 6a). These architectures with different combinations of hyperparameters were evaluated on the validation set. It is found that a ConvNet model had a less average L1 loss (0.1361) than transfer learning with a ResNet-50 (0.2370) or a MobileNet (0.2617) (Supplementary Figure 6b). This was probably because the ImageNet had very different image characteristics from our recordings. Another reason was that these computer vision models were developed to detect objects, but our goal was not to segment neurons but to extract embedded temporal information. Surprisingly, while batch normalization has been suggested to enhance model performance in the literature, we found our model without batch normalization had less L1 loss (0.1361) than using batch normalization (0.1450). We suspected that batch normalization removed fluctuations across samples, which might embed behavioral information, and thus compromised the decoding performance.

### 4.2 How behavioral states contributed to decoding errors

When comparing the decoding performance, we focused on periods when the animals were running at speeds above 5 cm/s (RUN), because previous studies have demonstrated that the hippocampal neurons had very different response profiles when the animal was resting or moved at very slow speeds (STOP) (Ahmed and Mehta, 2012; Davidson et al., 2009; Geisler et al., 2007;Ólafsdóttir et al., 2017). Indeed, when a trained ConvNet was evaluated on STOP periods, the decoded trajectories were very different from the true trajectories, as if the animal was mentally simulating running (Supplementary Figure 7a, b).

We also examined the neural manifolds of the decoder during STOP periods to evaluate how the maze topology was represented (Supplementary Figure 8a, b). Surprisingly, the maze topology was not completely lost during the STOP periods. Instead, a global maze topology was captured with fuzzy local structures, with reduced similarities (Last: 0.80 ± 0.02, stats =1, *p* < 0.001; Dense: 0.86 ± 0.02, stats = 5, *p* < 0.01) relative to manifolds during the RUN periods.

Nevertheless, these results demonstrated how internal states of the animal (e.g., whether the animal was running or not) could contribute to decoding errors. Whether we could use it as a tool to study hippocampal replays was beyond the scope of this study.

What contributed to decoding errors in the RUN periods? One common cause was the discrepancy between training and test sets. To evaluate the consistency of behavioral variables (*x, y, v*) between training and test sets, we compared the average occupancy and average speed in the space between the training and test sets (Supplementary Figure 1b and Supplementary Figure 9a), by computing their 2D correlations (*r*). Results showed that occupancy and average speed were mostly consistent throughout the session, but dropped near the end of the session (average *r* in T-maze: 0.71 ± 0.06, 0.74 ± 0.05, 0.78 ± 0.05, 0.76 ± 0.06, 0.71 ± 0.03; H-maze: 0.70 ± 0.03, 0.71 ± 0.02, 0.68 ± 0.06, 0.62 ± 0.04, 0.46 ± 0.13; □-maze: 0.75 ± 0.03, 0.75 ± 0.03, 0.72 ± 0.03, 0.66 ± 0.05, 0.70 ± 0.06).

We also quantified the similarity of each paired variable by computing structural similarity index measure (SSIM) between the training and test sets (Supplementary Figure 9b). If two joint distributions have the same structures, SSIM = 1. We found that the joint distributions in the T-maze experiments were highest, and joint distributions remained similar throughout the session in all datasets (average SSIM in T-maze: 0.82 ±0.03; H-maze: 0.72 ± 0.02; □-maze: 0.64 ± 0.04).

Overall, these analyses reveal a general consistency of behavioral variables across time, suggesting other factors contributing to decoding errors. One possibility was that the internal running-speed threshold varies across animals, such that some of the STOP periods were mistaken as the RUN periods, leading to decoding errors. If this was the case, we would observe a negative correlation between decoding errors and running speeds. Another possible source for decoding errors originated from limited temporal precision in calcium imaging. In this case, we would observe a positive correlation between decoding errors and running speeds. The other possible factor came from sampling bias in the training set. If there was a bias in the spatial occupancy, i.e., certain positions were visited more by the animal, we would observe a bias in predictions toward these frequently-sampled positions.

To evaluate these factors, where decoding errors occurred in the maze as well as where they pointed to were visualized (Supplementary Figure 10a), and compared with the average occupancy (x-y Density) and speed (AvgSpeed) maps in both training and test sets (Supplementary Figure 10b). Take a session from T-maze exploration as an example (Supplementary Figure 10a, b), where decoding error pointed to was mostly correlated with AvgSpeed map of the training set (*r* = 0.61), relative to the test set (*r* = 0.51) and x-y Density (training set: *r* = 0.38; test set: *r* = 0.34). On the contrary, where decoding error occurred was mostly correlated with AvgSpeed map of the test set (*r* =0.81), relative to the training set (*r* = 0.62) and x-y Density (training set: *r* = 0.50; test set: *r* = 0.52). This result was consistent across all the models (Supplementary Figure 10c). Where wrong predictions located were more correlated with the AvgSpeed map of the training set (*r* = 0.56 ± 0.02; stats = 134, *p* < 0.05), whereas where error occurred was mostly correlated with the AvgSpeed map of the test set (*r* = 0.78 ± 0.01, blue; Wilcoxon signed-rank test: stats =1, *p* < 1*e* − 5)

### 4.3 Information embedded in background residuals

Our findings demonstrate that behavioral variables such as position and speed can be directly decoded from raw microendoscopic images, without a need to specifically identify putative neurons and/or deconvolve spikes. Our ConvNet-based approach outperforms a feedforward neural network that takes the denoised calcium traces as input, suggesting that neuropil/background, which is filtered in a typical calcium analysis pipeline, encodes behavioral information. One might argue that this performance difference is driven by different decoder architectures or input dimensions. However, by generating images composed of isolated background and foreground components as input for the ConvNet decoder, we showed that the residual signals encode the animal’s position and speed. By occluding spatial footprints of the neurons, with mostly neuropil signals remained, we excluded the possibility of successful decoding being a byproduct of incomplete source extraction within the ROIs. Our results indicate that neuropil/background in single-photon calcium imaging in the hippocampal CA1 contains spatial information.

Neuropil signals are considered to be out-of-focus fluorescence signals from nearby cellular processes including dendritic spines and axonal segments (Gobel and Helmchen, 2007; Kerr et al.,2005; Ohki et al., 2005), and are often removed in a calcium imaging analysis pipeline (Keemink et al., 2018; Lu et al., 2018; Pachitariu et al., 2017; Pnevmatikakis et al., 2016; Zhou et al., 2018). However, studies have shown that dendritic fluorescence can exhibit calcium transients in response to synaptic inputs, back-propagating action potentials and dendritic spikes (Kleindienst et al., 2011; Larkum et al., 1999; Svoboda et al., 1999; Takahashi et al., 2012; Yuste and Denk, 1995), which are important for information processing. For example, dendritic spikes in the sensory cortex can contribute to the orientation selectivity (Euler et al., 2002; Sivyer and Williams, 2013; Smith et al., 2013) and angular tuning (Lavzin et al., 2012). In addition, dendritic signals in the CA1 can modulate the place fields and induce novel place field formations to produce feature selectivity (Bittner et al., 2015; Sheffield and Dombeck, 2015).

Similarly, axonal calcium imaging has shown that individual axons can encode task-related variables such as touch and whisking (Petreanu et al., 2012). Together, these suggest that neuropil signals, comprising both dendritic and axonal activities, encode external stimuli and behavioral information. This hypothesis is supported by our results, consistent with previous studies that demonstrate both cell somata and neuropil patches can represent goal-directed behaviors (Allen et al., 2017) as well as orientations of the moving gratings (Lee et al., 2017). To isolate the contributions from dendrites, axons, and out-of-focus cell somata in the background signals, a calcium indicator which specifically localizes to the soma (e.g., SomaGCaMP (Shemesh et al., 2020)) or the axon (e.g., axon-GCaMP6 (Broussard et al., 2018)) will be needed for future experiments. Overall, our study challenges the idea of removing background signals from the calcium imaging data, and proposes to re-examine the analysis pipeline for extracting behavioral information from microendoscopic data.

### 4.4 Contributions from our end-to-end decoding paradigm

In addition to uncovering information embedded in imaging background, our work does not need sophisticated hyperparameter tuning that requires prior knowledge, and provides an easy-to-implement framework for decoding behavior from imaging data. Many methods have been proposed to decode position from activities of a group of single neurons in the hippocampus in the literature, but they require either spike sorting for electrophysiology recordings (Brown et al.,1998; Wilson and McNaughton, 1993; Zhang et al., 1998) or spike inference for calcium imaging data (Etter et al., 2020; Gonzalez et al., 2019; Shuman et al., 2020; Stefanini et al., 2020; Ziv et al., 2013). Although alternative decoding methods for calcium imaging have been developed to bypass spike deconvolution, such as estimating neuronal firing rates directly (Ganmor et al., 2016), approximating spiking with marked point process (Tu et al., 2020), or decoding using frequency representations (Frey et al., 2019), they all require extracting fluorescence traces from identified individual neurons. On the other hand, new statistical methods are developed to decode position with unsorted spikes or local field potentials in electrophysiology (Cao et al., 2019; Deng et al.,2015; Frey et al., 2019; Kloosterman et al., 2014). However, this idea has never been applied in single-photon calcium imaging. This study is the first demonstration to directly decode behaviors from raw microendoscopic data without the need to demix individual neuronal signals.

Our approach benefits from the power of deep learning. With recent advances in machine learning, there have been efforts to use deep learning models such as a recurrent neural network to learn hidden features from neural data and decode animal behavior with high accuracies (Glaser et al.,2020; Tampuu et al., 2019). However, these efforts still rely on the identification of single neurons and unfortunately, fail to take full advantage of deep learning methods to extract information from noisy data. Our end-to-end decoder embeds all preprocessing steps into the network. In addition, we adapted the Grad-CAM algorithm (Selvaraju et al., 2017) to find saliency maps across layers of the decoder. We chose Grad-CAM given its reduced visual artifacts (Adebayo et al., 2018) and smoother saliency maps compared to other approaches such as Vanilla Gradients (Simonyan et al., 2013), Guided Backpropagation (Springenberg et al., 2014) and Deconvolution (Zeiler and Fergus, 2014). However, this method does have a limitation for capturing fine-grained detail. Nevertheless, our results showed that video decomposition automatically emerges in a trained decoder, and identified clusters composed of different groups of cells and neuropil patches for spatial representations in the CA1. These results offer an alternative way to classify neural activities by visualizing the saliency maps of the decoder, without a need to run an extra clustering algorithm.

In conclusion, we have (1) demonstrated an efficient and easy-to-implement decoder for raw microendoscopic data, (2) revealed extra behavioral representations embedded in neuropil background, and (3) proposed a method to identify clusters relevant for extracting position and speed information in neural data. We believe that our decoding analysis can be extended to examine representations of other brain regions such as the entorhinal cortex, primary visual cortex, and parietal cortex, underlying different behavioral tasks, with potential applications in real-time decoding and closed-loop control.

## Author Contributions

C.-J.C. and M.A.W. conceived the project; C.-J.C. developed models and experiments, analyzed the data, and wrote the code; W.G. and J.J.Z. provided animal behavioral and calcium imaging data; J.P.N. developed techniques for data collection; W.G. implemented the CNMFe algorithm and labeled the data; S.-H.S. provided consultation for Tensorflow infrastructure; C.-J.C prepared the manuscript; M.A.W. supervised the project. All authors contributed to manuscript editing.

## Supplementary Figures

**Supplementary Figure 1:**
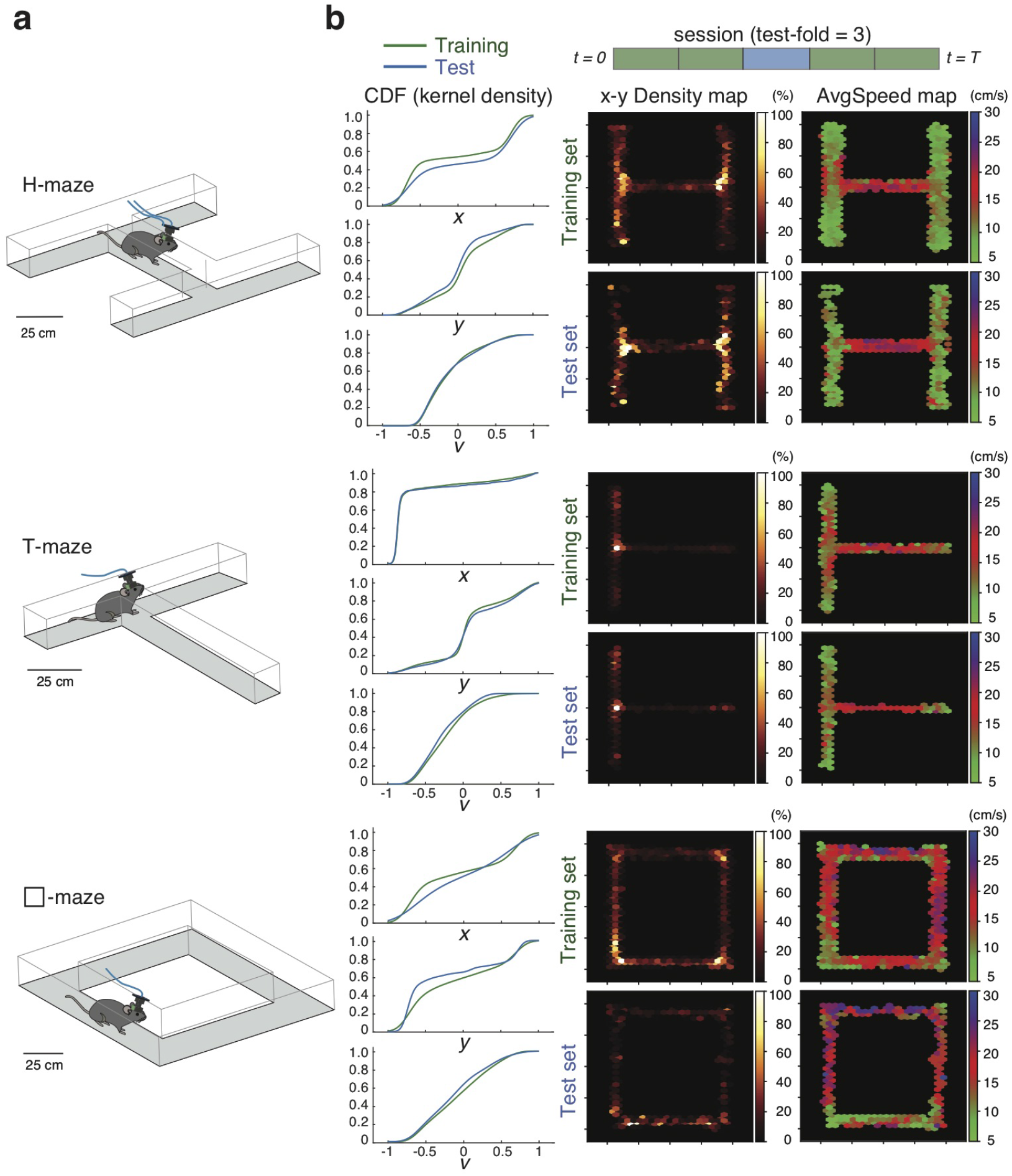
Examples of H-maze, T-maze and □-maze exploration behavior. (**a**) Schematic diagrams for H-maze (*top*), T-maze (*middle*) and □-maze (*bottom*) exploration experiments. (**b**) Evaluation of a train/test split for each maze exploration. Left: The cumulative density function (CDF) of each label between training and test sets. Right: Average occupancy (x-y Density) and speed maps (AvgSpeed) for a typical train/test split. For an example H-maze, T-maze, and □-maze dataset, its 2D correlation between training and test sets: r(x-y Density) = 0.7230, 0.9154, and 0.5232, respectively; r(AvgSpeed) = 0.7678, 0.7379, and 0.6612, respectively.

**Supplementary Figure 2:**
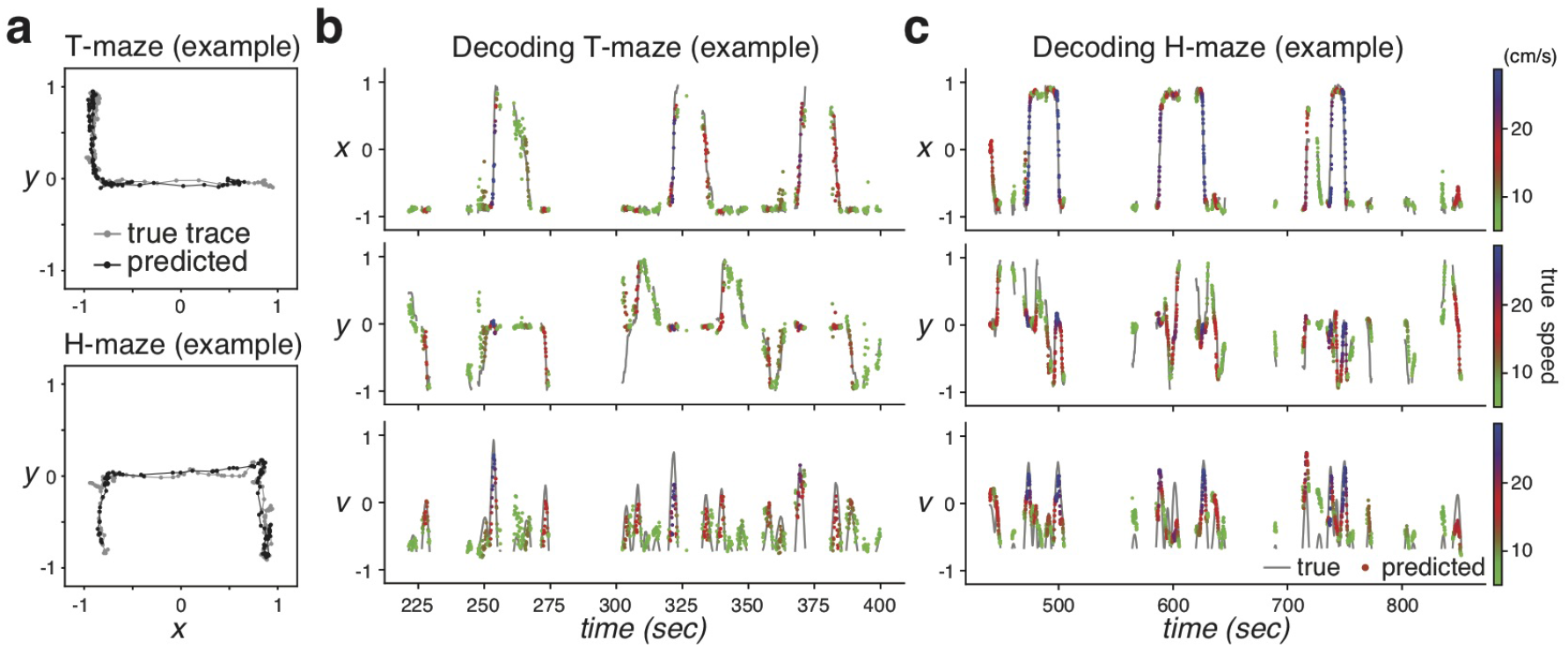
Examples of decoded running traces in the T-maze and the H-maze. (**a**) Top: Example 10-sec trajectories from an animal in a T-maze exploration experiment. Bottom: Example 10-sec trajectories from an animal in a H-maze exploration experiment. (**b**) Decoded variables across time in an example T-maze test set. Average error in decoding x, y, and v: 2.95 cm, 6.15 cm, 3.12 cm/s, respectively. (**c**) Decoded variables across time in an example H-maze test set. Average error in decoding x, y, and v: 5.53 cm, 9.94 cm, 3.65cm/s, respectively.

**Supplementary Figure 3:**
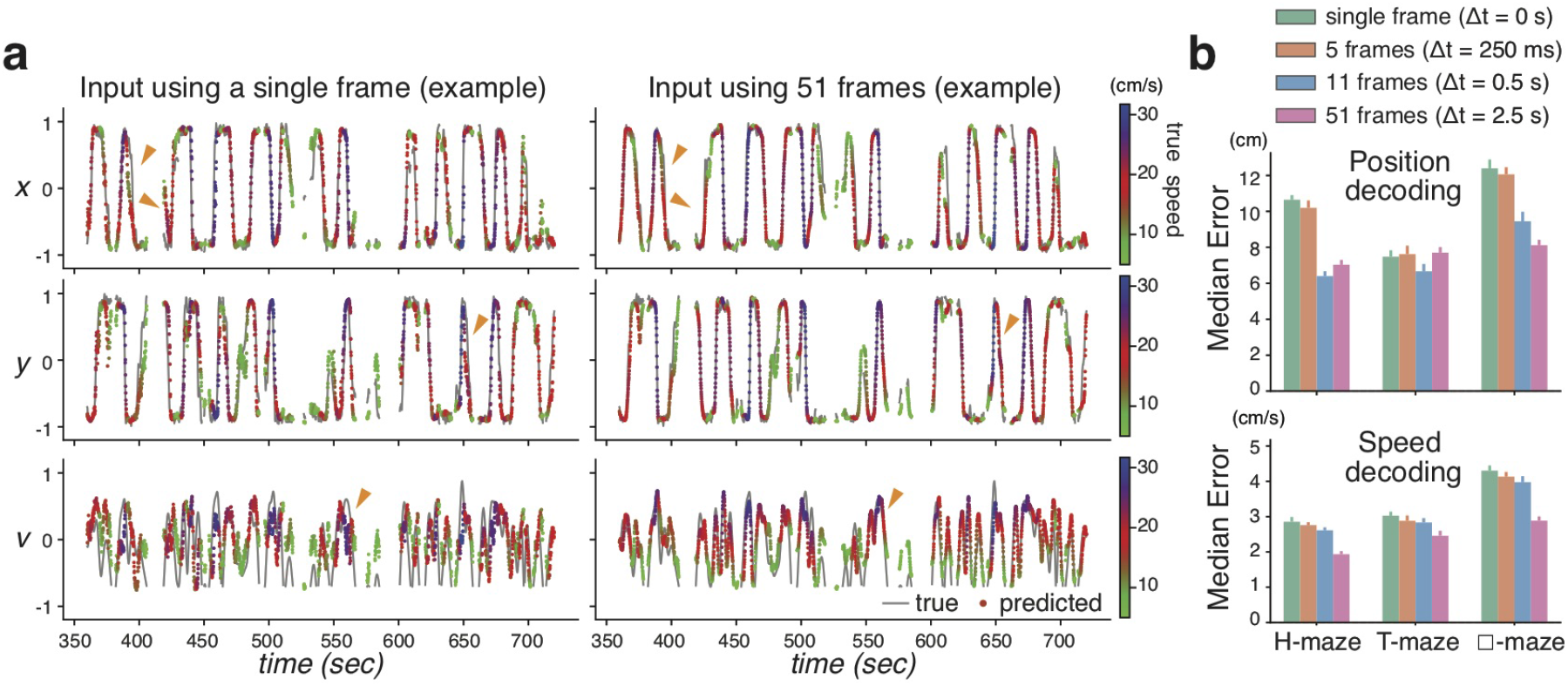
Optimal decoding using a 1-second input and convolutions independent of time. (**a**) Examples of variables decoded from the same dataset, using a single-frame input (*left*) and 51 frames (*right*) when convolutions were independent of time. Orange triangles mark timepoints where we can easily see prediction differences between models. The gray line denotes true traces. Dots denote predicted traces, colored based on running speeds. (**b**) Model performance across all datasets. Top: Incorporating 11 frames (time window of 1 s) led to least median position-decoding error (7.50 ± 0.60 cm), significantly different from single frame input (10.17 ± 0.43 cm; stats = 5, *p* < 1*e* − 5). Bottom: Incorporating 51 frames (time window of 5 s) led to least median speed-decoding error (2.42 ± 0.09 cm/s), significantly different from single frame input (3.39 ± 0.14 cm/s; stats = 1, *p* < 1*e* − 5).

**Supplementary Figure 4:**
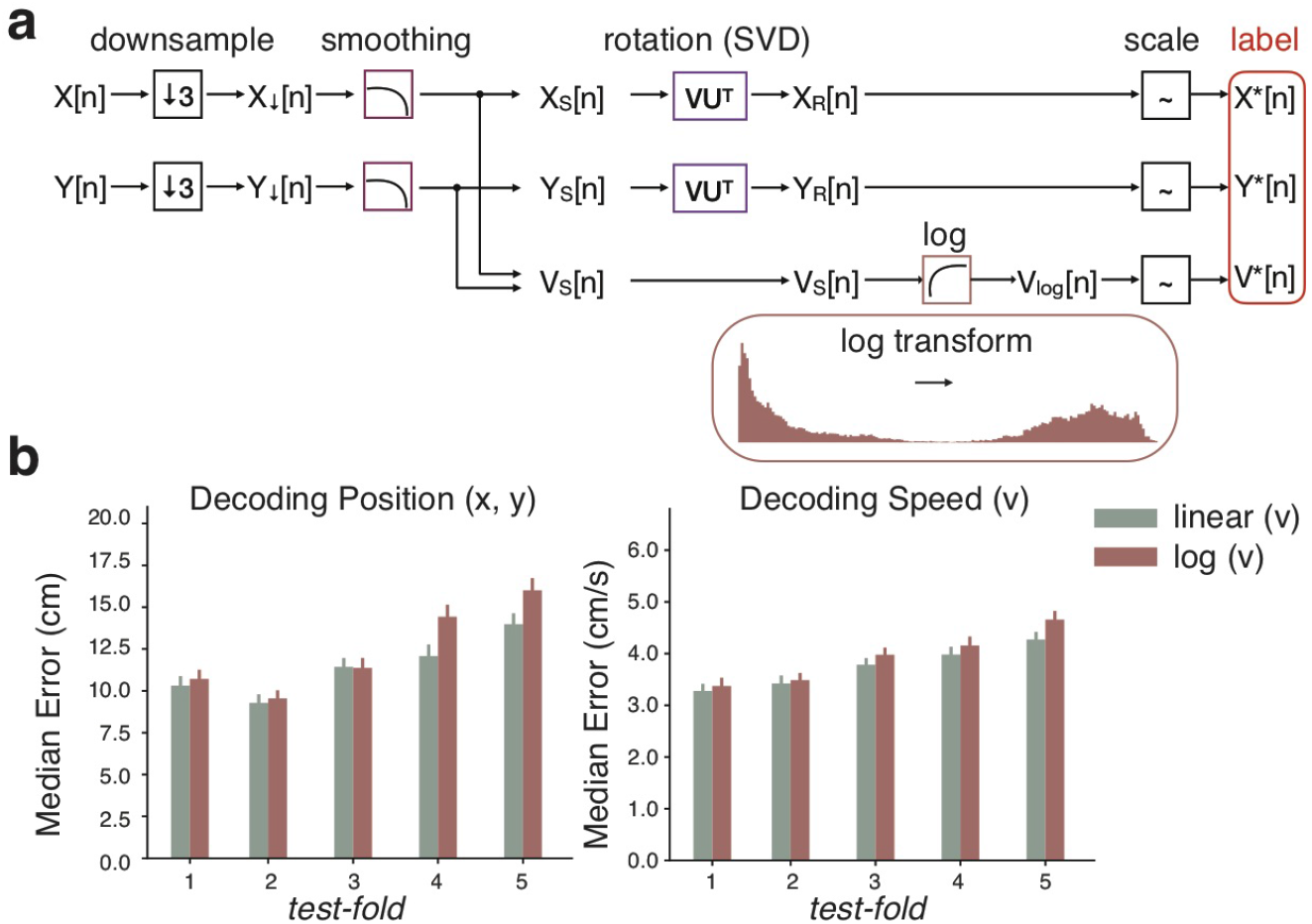
Decoding performance with log transformation on running speeds. (**a**) Schematic diagram of data preparation with speeds being log-transformed to create a log-uniform distribution. (**b**) Model performance across a subset of datasets across different test-folds. Left: The original model trained with linear speeds (11.49 ± 0.13 cm, sage) had significantly less median error in predicting positions than the same model trained with log speeds (12.45 ±0.14 cm, cardinal; Wilcoxon signed-rank test: stats = 15274374, *p* < 1*e* − 5). Right: The original model trained with linear speeds (3.76 ± 0.03 cm/s, sage) had significantly less median error in predicting speeds than the same model trained with log speeds (3.95 ± 0.04 cm/s, cardinal; Wilcoxon signed-rank test: stats = 15506009, *p* < 1*e* − 5).

**Supplementary Figure 5:**
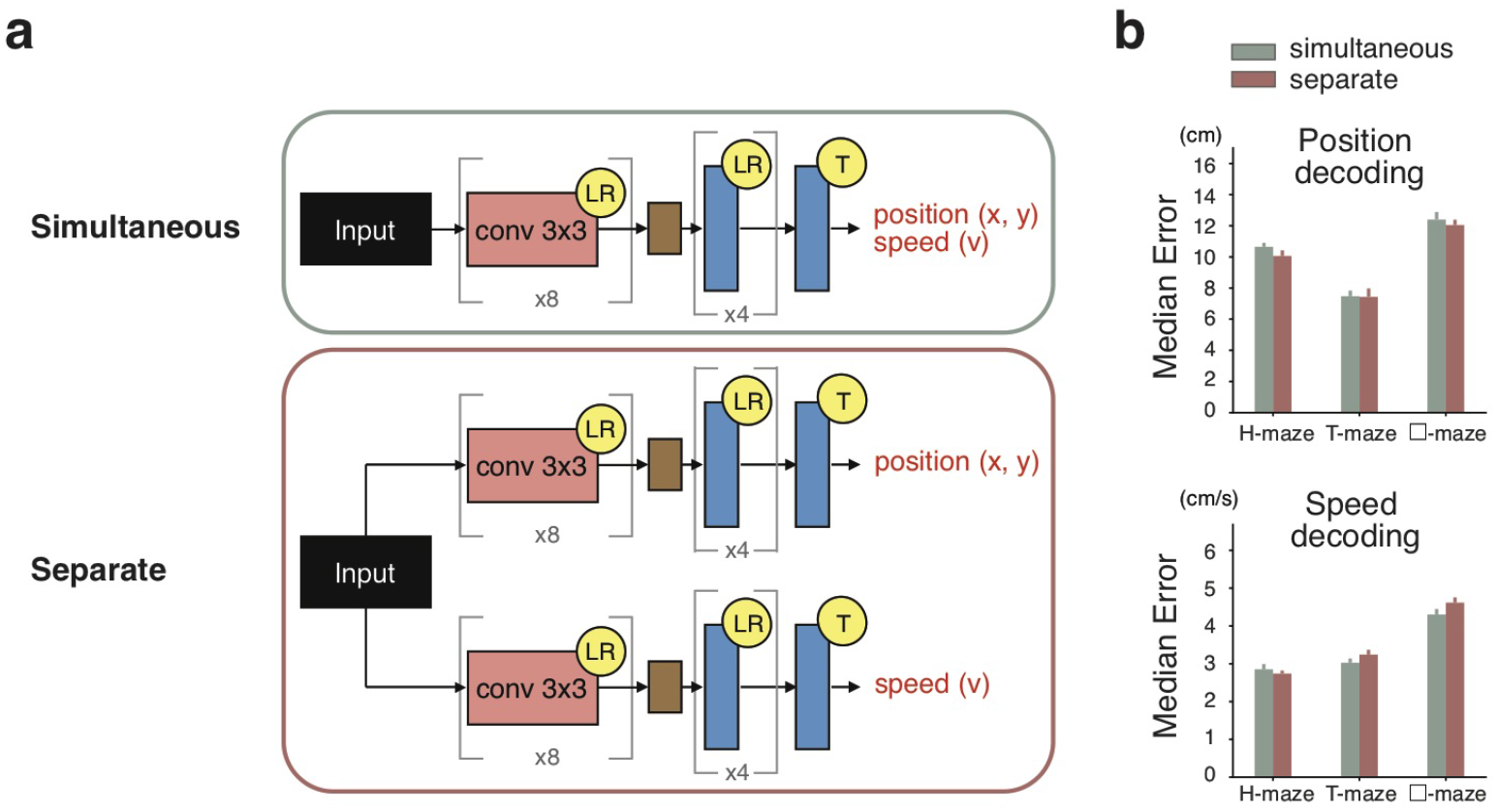
Network performance with separate streams for decoding behavior. (**a**) Schematic diagram of ConvNet architectures with original simultaneous (top) and separate (bottom) decoding strategies. (**b**) Model performance across all datasets. Top: The original model with simultaneous decoding (10.17 ± 0.43 cm, sage) had significantly more median error in predicting positions than the model with separate decoding streams (9.85 ± 0.42 cm, cardinal; Wilcoxon signed-rank test: stats = 133, *p* < 0.05). Bottom: The original model (3.39 ± 0.14 cm/s, sage) had significantly less median error in predicting speeds than the model with separate decoding streams (3.53 ± 0.16 cm/s, cardinal; Wilcoxon signed-rank test: stats = 131, *p* < 0.05).

**Supplementary Figure 6:**
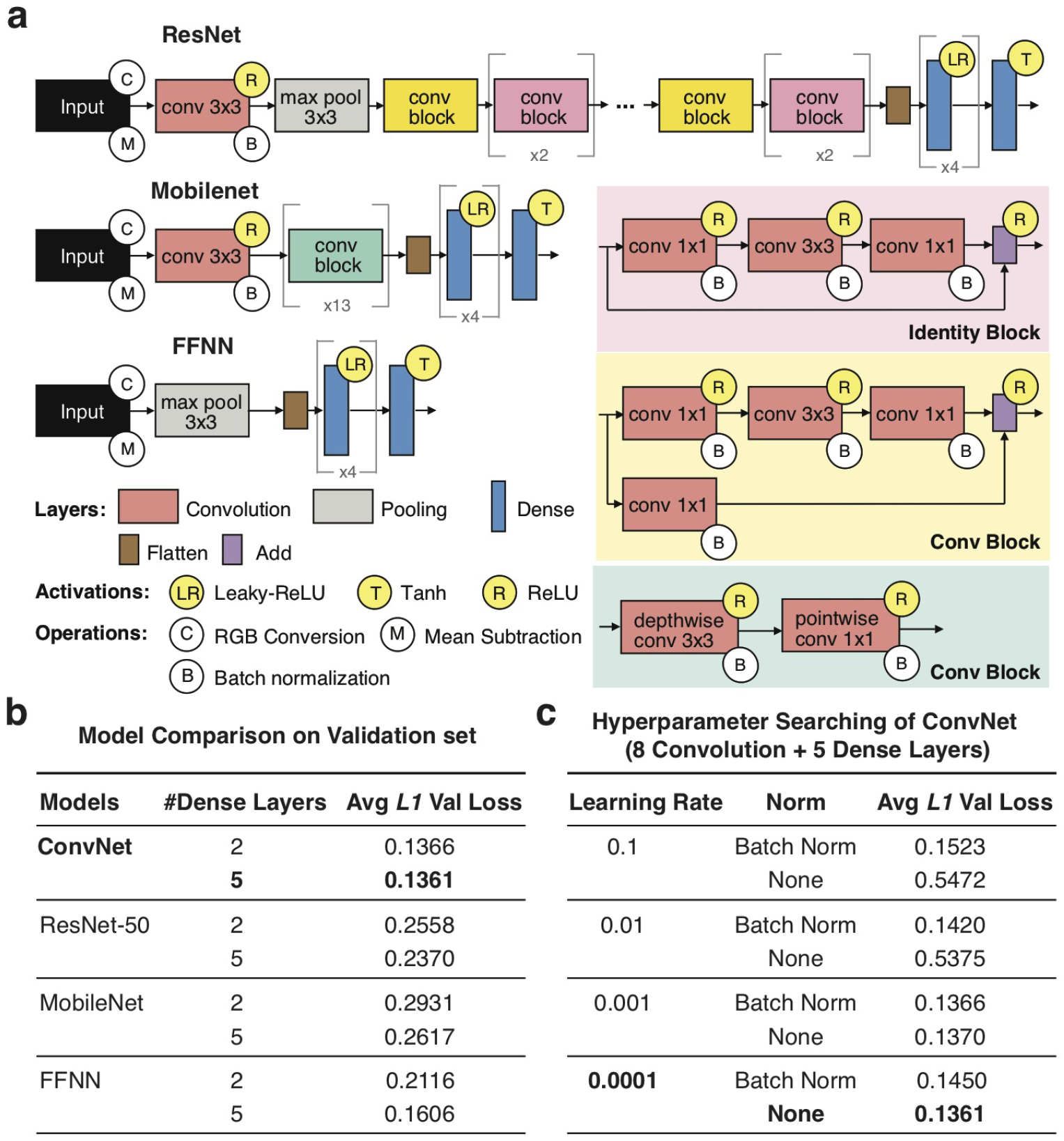
Model selection with hyperparameter searching. (**a**) Schematic diagram of different model architectures. We did transfer learning with a ResNet model (top) and a Mobilenet model (middle) pre-trained on Imagenet. We also designed a feedforward neural network, where max pooling was applied on input images before feeding flattened features to the dense layers. Hyperparamter searching was applied to all models and the best combination was selected based on validation performance. Details see Materials and Methods. (**b**) Model comparison on validation set. We compared all models with their best hyperparameter sets, and found the ConvNet model had the least average L1 loss. (**c**) Subsets of hyperaparameter searching on validation set. For a ConvNet model with 8 convolution with 8 convolutional layers and 5 dense layers, we found training model with learning rate at 1e-4 without batch normalization had least average L1 loss. These hyperparameters were used for models in all the experiments.

**Supplementary Figure 7:**
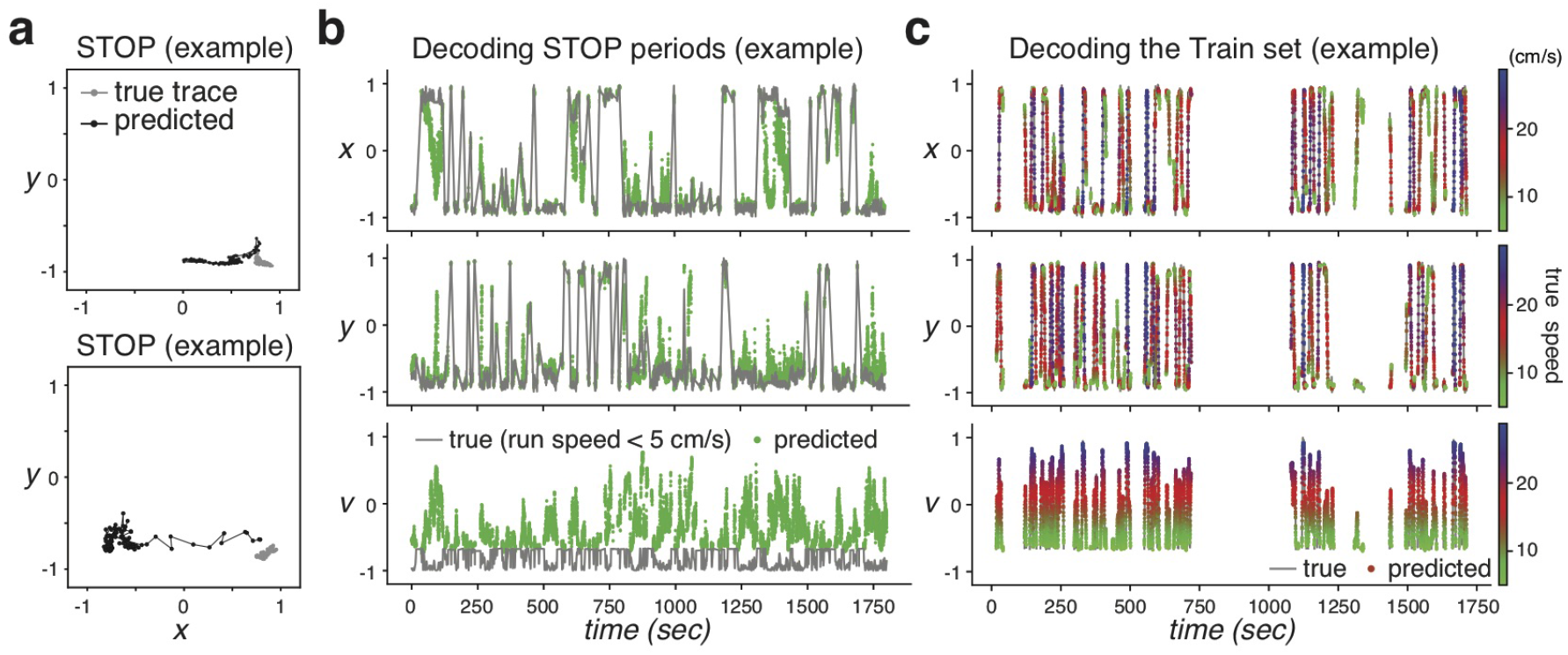
Examples of decoded traces in STOP periods and the training set. (**a**) Example 10-sec trajectories of true (gray) and decoded (black) positions when the animal was mostly stationary. Decoded traces formed a sequence as if the animal was simulating running. (**b**) Decoded variables across time during STOP periods in an example test set. Gray line denotes true traces. Green dots denote predicted traces. (**c**) Decoded variables across time in an example training set. Dots denote predicted traces, colored in actual running speeds.

**Supplementary Figure 8:**
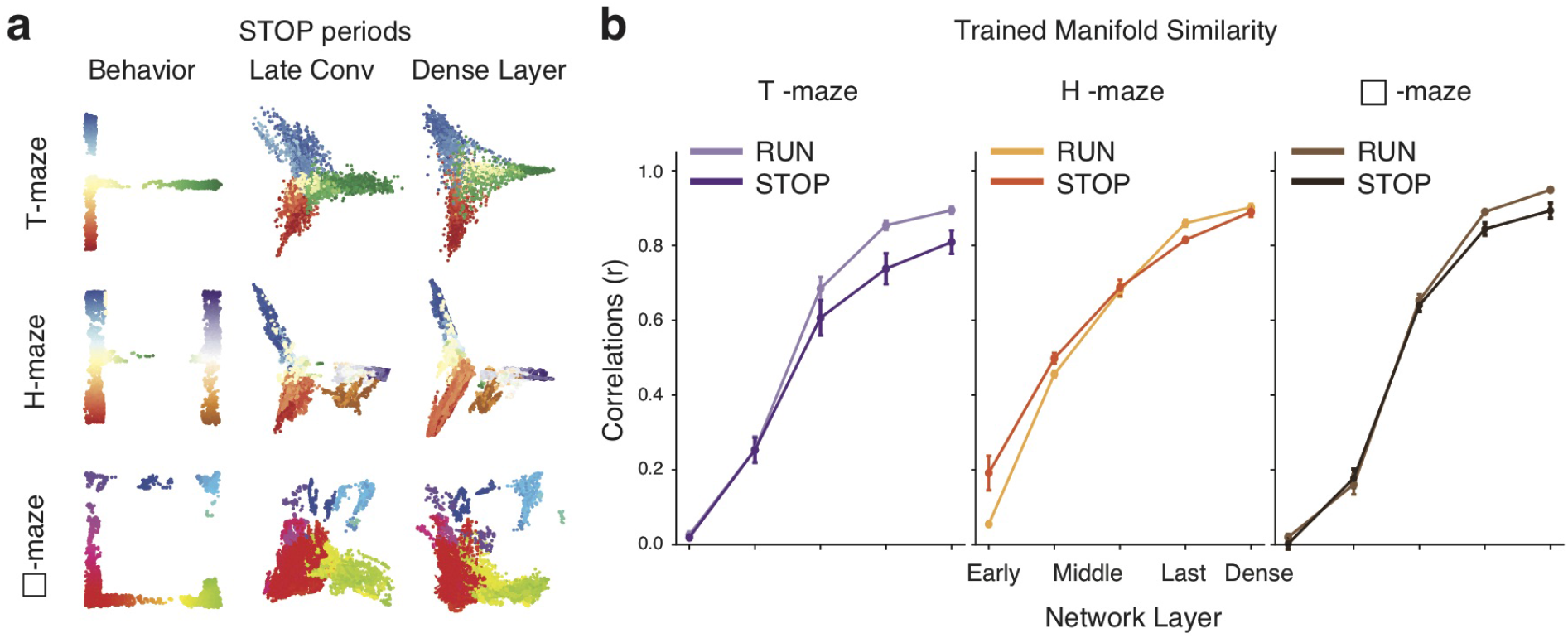
Encoding of behavioral topology during STOP periods in manifolds across model layers. (**a**) Ensemble representations from the last convolutional layer and the dense layer of a trained model on a typical dataset in the T-maze (*top*), H-maze (*middle*), and □-maze (*bottom*) (same models as in **Figure 7a**). (**b**) Comparison between neural manifolds extracted from RUN and STOP periods (i.e., animals were either stationary or running at speeds below 5 cm/s), in the T-maze (*left*), H-maze (*middle*), and □-maze (right). Across all datasets, manifold similarity was significantly higher when the animals were running (Wilcoxon signed-rank test: stats = 1, *p* < 0.05).

**Supplementary Figure 9:**
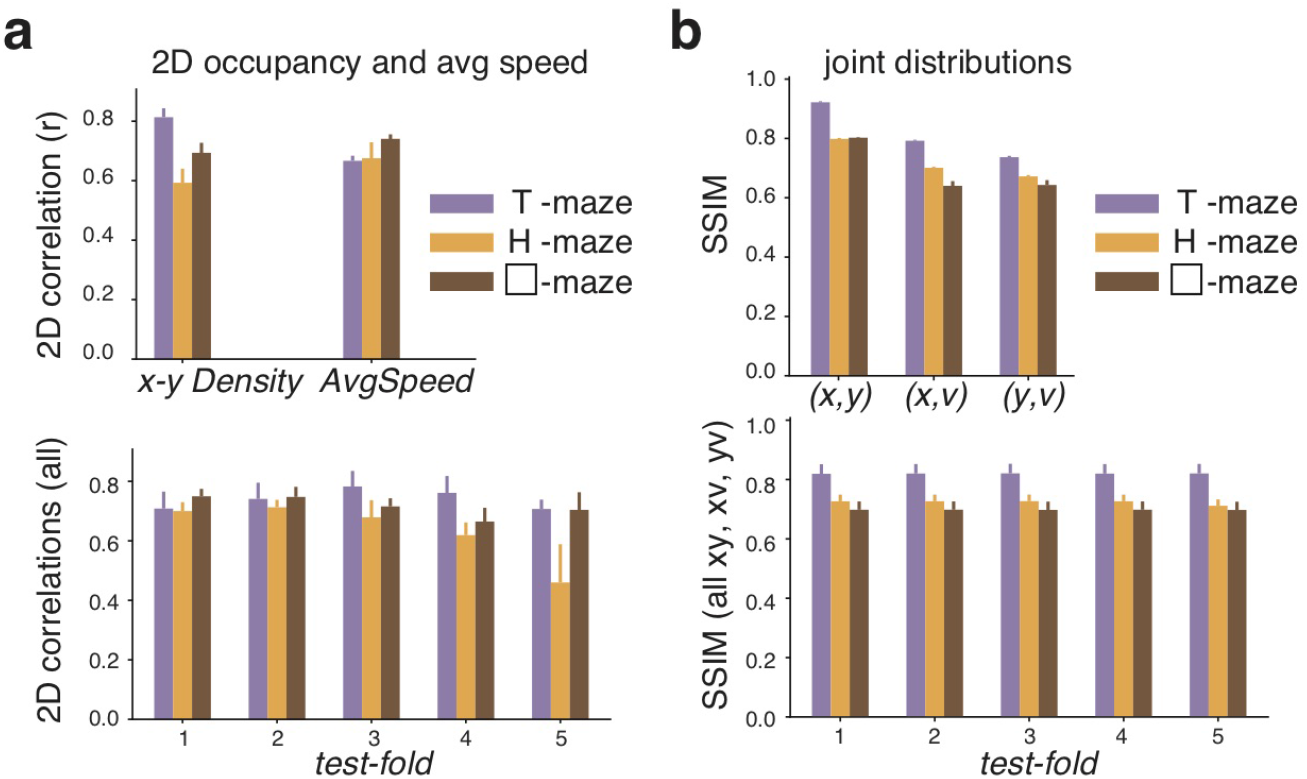
Consistency of behavioral distributions between training and test sets. (**a**) Similarity of average occupancy (x-y Density) and average speed (AvgSpeed) between the training and test sets. Top: 2D correlations of occupancy and speed maps in different mazes. Bottom: All 2D correlations (x-y Density, AvgSpeed) across test-folds. (**b**) Evaluation of train/test split using structural similarity index measures (SSIM) on joint distributions of output labels. Top: SSIM for each paired variables (*x, y*), (*x, v*) and (*y, v*) in different maze experiments. Bottom: SSIM ((*x, y*), (*x, v*) and (*y, v*)) across test-folds in different maze experiments.

**Supplementary Figure 10:**
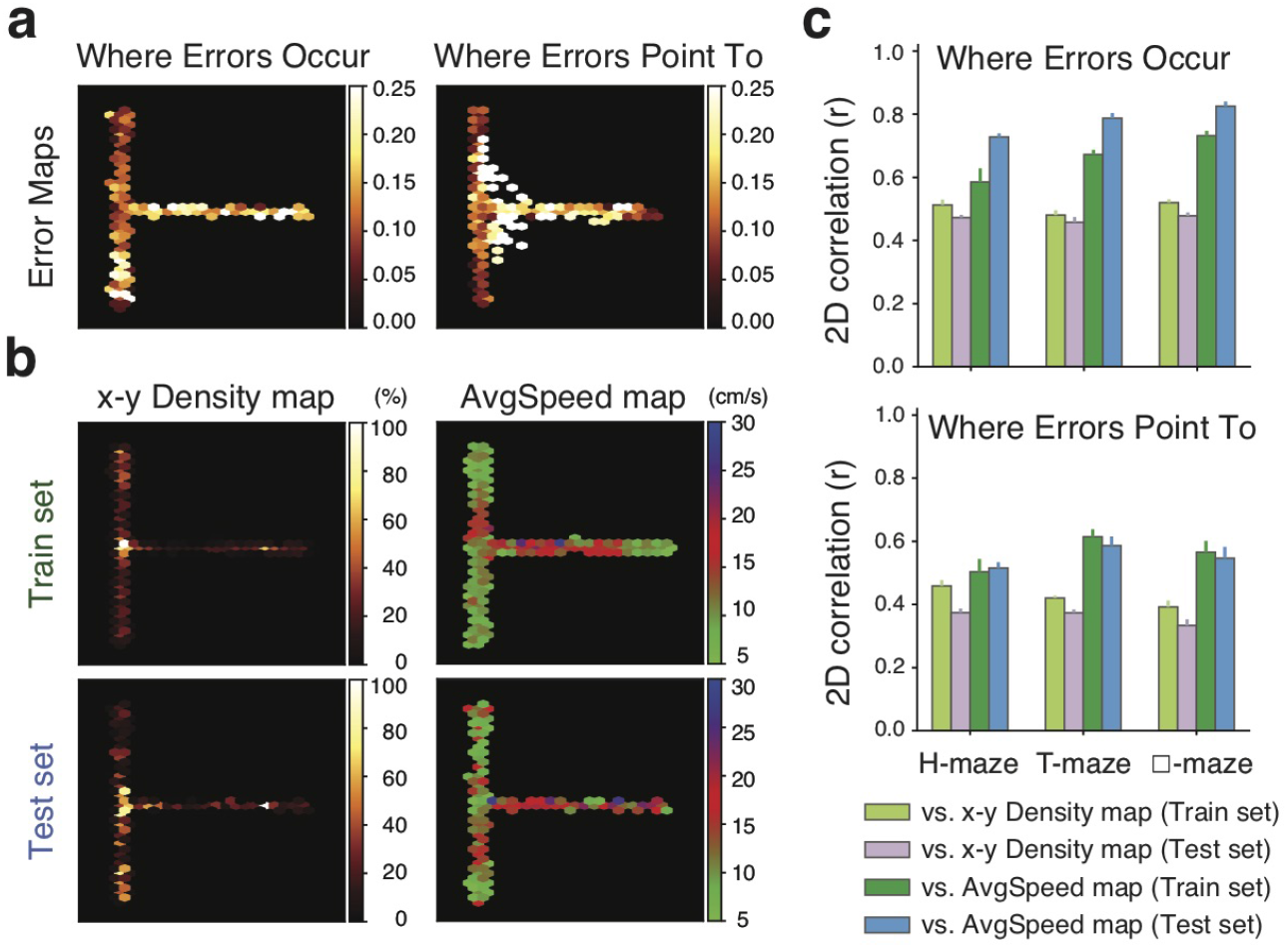
Possible factors contributing to decoded errors. (**a**) Error visualizations in the space from an example T-maze exploration dataset. Heatmaps represent decoded errors (a.u.) before converting back to the original scales (cm and cm/s). Left: Spatial maps of where decoded errors occurred. Right: Spatial maps of where errors pointed to (i.e., where wrong predictions were). (**b**) Left: Average occupancy (x-y Density) maps of the training set *top* and the test set *bottom* from the example dataset in **a**. Right: Average speed (AvgSpeed) maps of the training set *top* and the test set *bottom* from the example dataset in **a**. 2D correlations (*r*) between these maps to the error maps were computed to evaluate potential contributing factors for decoded errors. Here, where error pointed to was mostly correlated with the AvgSpeed map of the training set (*r* = 0.61, right, top), and where error occurred was mostly correlated with the AvgSpeed map of the test set (*r* = 0.81, right, *bottom*). (**c**) Across all datasets, where error pointed to was more correlated with the AvgSpeed map of the training set (*r* = 0.56 ± 0.02, green; Wilcoxon signed-rank test: stats = 134, *p* < 0.05) relative to other factors, but overall the correlations were mild. On the other hand, where error occurred was mostly correlated with the AvgSpeed map of the test set (*r* = 0.78 ± 0.01, blue; Wilcoxon signed-rank test: stats = 1, *p* < 1*e* − 5)

## 5 Materials and Methods

### 5.1 Data collection

A total of 8 male mice of C57/B6 background were used in the experiments. All procedures were approved by the MIT Committee on Animal Care and followed the guidelines established by the National Institutes of Health for the care and use of laboratory animals.

#### Virus-mediated gene delivery

Animals were first anesthetized with 5% Isoflurane in oxygen. Throughout the procedure, the animals were anesthetized at a stereotaxic surgical frame (Kopf) with continuous 1.5% to 2% Isoflurane in oxygen. A burr hole is made at −1.5mm medial-lateral and 2.1mm rostral-caudal to the Bregma landmark. Using a motorized injector at a speed of 0.05 *μ*L/min, 0.5*μ*L of pGP-AAV1-syn-jGCaMP7f-WPRE (titer: 1.9 x 1012 gc/ml) was injected at 1.5mm under the skull surface in the burr hole. Afterwards, the scalp was sutured and analgesic was administered (Buprenex, 0.05 mg/kg) prior to the animals’ recovery.

#### Implant protocol

After virus injection, GRIN lens implant and baseplate implant were performed separately. During implant, the animals were anesthetized following the same protocol.

##### GRIN lens

The skull surface was roughened with etchant gel (C&B Metabond). A craniotomy 2 mm to 2.2 mm in diameter was made on the skull, with the burr hole from virus injection procedure in the center. Cortical tissue was aspirated out, exposing the rostral-caudal running striation pattern right above CA1. A GRIN lens (1.8 mm diameter, 0.25 pitch, 670 nm, Edmund) was perpendicularly placed on top of CA1 and fixed to the skull with adhesive (Loctite 454). Afterwards, the exposed skull and the surrounding of the GRIN lens were covered with transparent dental cement (C&B Metabond), followed by black orthodontic acrylic resin to block out ambient light. The top of the GRIN lens was covered with silicone adhesive (Kwik-Sil).

##### Baseplate

With a baseplate attached to a microendoscope (UCLA miniscope), the optimal field of view (FOV) was searched through imaging and manually adjusting the angle of the baseplate attachment. After the optimal field of view was reached, the baseplate was first fixed in place with adhesive (Loctite 454), and then affixed to the skull with dental cement (M&B metabond).

#### Imaging setup

One-photon calcium imaging was performed through a head-mounted miniscope. The imaging power was between 1 mW and 10 mW approximately, adjusted for each animal. The excitation LED is filtered by a 470/40 band-pass filter. The emitted photons were collected by a CMOS sensor, filtered by a 525/50 band-pass filter, sampled at 30 Hz. The imaging FOV was 480 by 752 pixels, and the optimal plane was reached during the baseplate implant.

### 5.2 Behavioral Tasks

Once the animals were fully recovered from previous procedures (approximately 3 days after the baseplate implant), *in vivo* recordings were performed on awake animals.

#### Miniscope acclimation in the home cage

The animals were attached to the imaging cables, allowing them to acclimate to the miniscope and tethering for a few sessions in the home cage, with each session lasting for 30 minutes. This procedure repeated for about 3 days, until the animals were comfortable with the imaging setup and exhibit normal behavior.

#### Maze exploration experiments

After miniscope acclimation sessions, the animals underwent maze exploration experiments in which maze enclosures are made of corrugated plastic sheets (Home Depot). The shapes of these mazes can be categorized as T-shape, H-shape, or □-shape. Each maze has a size of about 1 m by 1 m. For each maze, the animals underwent multiple sessions, with each session lasting for about 30 minutes. In our study, we excluded the first few sessions when the maze configuration was still novel to the animals, and only used data in later sessions when the animals were already familiar with the environment.

### 5.3 Dataset preparation

#### Behavioral data

To track the animals’ behavior, we used a webcam sampled at 60 Hz and synchronized with imaging using a package (https://github.com/jonnew/Bonsai.Miniscope) for Bonsai (Lopes et al., 2015). Red tape on the head-mounted miniscope was tracked as a proxy for the animal’s locations, reported as position coordinates (*x, y*).

##### Temporal down-sampling and smoothing

Raw x, y coordinates were downsampled to 10 Hz, and smoothed with a Savitzky-Golay finite impulse response smoothing filter of quadratic order. The speed *v* was further estimated from these smoothed x, y coordinates.

##### Coordinate orthogonalization

Linear dependencies between different dimensions of the output labels were minimized by orthogonalizing the smoothed x, y coordinates. For the T-maze, x, y coordinates were centered at the intersection of two perpendicular arms. For the H-maze and the □-maze, x, y coordinates were centered in the middle of the maze. After centering, a rigid rotation matrix (R) was estimated through singular value decomposition (SVD) and applied to orthogonalize the x, y coordinates.

##### Label scaling

All behavioral labels (*x, y, v*) were scaled to the range [-1, 1].

#### Calcium Imaging

##### Movement artifact removal

To correct non-uniform movement artifacts caused by brain motion, NoRMCorre, a fast non-rigid registration method (Pnevmatikakis and Giovannucci, 2017) was applied. This algorithm operates by splitting the FOV into overlapping spatial patches, which are further registered at a subpixel resolution for rigid translation against a template. Afterwards, a smoothed motion field for each frame was created by up-sampling the estimated alignments.

##### Spatial down-sampling and cropping

Images were down-sampled in each dimension by a factor of 2, leading to 240 by 376 pixels. The FOV was cropped to 224 by 224 pixels, by removing surrounding regions that did not contain calcium signals.

##### Source decomposition and denoising

This step was only applied to generate data for the baseline decoder (i.e., it was skipped for our end-to-end decoder). To separate different sources such as single neurons and out-of-focus fluorescence from neuropil, the CNMFe algorithm (Zhou et al.,2018) was applied. Hyperparameters such as the estimated diameter of a neuron were optimized for each animal’s dataset. The average fluorescence traces C from regions of interest (ROIs) were extracted and denoised through spike deconvolution. ROIs with weak (peak C lower than 100% dF/F) or sporadic activity (less than 3 calcium transients) were eliminated. The remaining ROIs were assumed to be putative neurons. Temporal activities C were used as input for the baseline decoder.

##### Temporal down-sampling

Calcium images were down-sampled to 10 Hz.

##### Pixel scaling

Raw image pixel amplitudes (0-255) were scaled to the range [-1, 1].

#### Experiment on residual information

##### Raw

This set consisted original images (*Y*) after movement artifact removal, FOV cropping, and pixel scaling.

##### Clean

A video matrix was obtained after the CNMFe algorithm by multiplying the spatial footprint matrix *A* with the temporal dynamics matrix *C*. Image frames from this video matrix only contained putative neurons, and thus were called Clean images.

##### Residual

Residual images were obtained by subtracting the Clean images from the Raw images, i.e. *Y* − *AC*. Theoretically, behavioral-relevant information in these images came from the background residuals, as any spike-triggered temporal fluctuations in the putative neurons were extracted to the temporal dynamics matrix *C*. Ideally, Residual images lack information from putative neurons. However, the CNMFe algorithm updates parameters through alternating iterations such that spike-triggered temporal fluctuations might not be fully captured in the Clean images. To further rule out spike-triggered information left in the Residual images, locations at putative neurons were occluded, through either Hollow ROI or Hollow A methods.

##### Hollow ROI

The Hollow ROI images were obtained after occluding the putative cells after setting a local adaptive threshold. The threshold was computed for each pixel according to the image characteristics within a moving circular window. The local threshold was computed using the Niblack method:

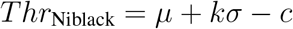

where k = 0.2, c = 0, *μ* is the mean pixel value within the window, and *σ* is the standard deviation of the pixel values within the window.

##### Hollow A

The Hollow A images were obtained after occluding the entire spatial footprint matrix A from the Residual images. This further removed any local signals associated with each cell.

##### Scramble images

Each pixel in each image was randomly re-positioned per row. The pixel distributions remained identical, but the spatial patterns were destroyed.

##### Random images

Image frames were synthesized by sampling each pixel from a uniform distribution that had the same minimum and maximum as the original image frames. The synthesized images were random noise. Decoders trained on random images were used as the chance level.

### 5.4 Decoder Modeling

#### Dataset evaluation

##### Train/val/test dataset split

Dataset was separated into the ‘RUN’ and the ‘STOP’ periods based on running speed with a separation threshold at 5 cm/s. For most of our model training and evaluation, the ‘STOP’ period was excluded from analysis unless otherwise indicated. Each animal’s dataset was segmented into 5 consecutive blocks. Following a 80-20 split rule, one of these blocks was selected to be the test set whereas the rest were used as the training set. During the model development phase, the training set was segmented into 5 consecutive blocks, with one of them randomly selected to be the validation set.

##### Similarity metrics for joint distribution

To quantify similarity between joint distribution for paired variables ((*x, y*), (*x, v*), (*y, v*)) between the training set and the test set, structural similarity index measure (SSIM) (Wang et al., 2004) was performed with *scikit-image* library. The joint distribution of the training set was viewed as an image (Im_train_), and so was the joint distribution of the test set (Im_test_). Each image was convolved with a normalized Gaussian kernel of width *σ* = 1.5, resulting in multiple patches. SSIM was calculated based on various patches from two images.

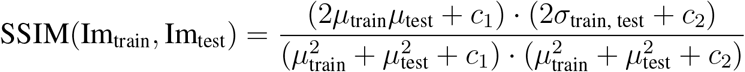

where *c*_1_ and *c*_2_ are constant, *μ* and *σ* are statistical attributes of all these patches. If SSIM = 1, two images of joint distributions have the same textures. A lower SSIM suggests more differences between two sets.

##### Similarity metrics for average occupancy map and average speed map

The occupancy map, i.e., x-y Density map, is a 2D hexagonal binning plot quantifying normalized occupancy on the maze. Average speed map, i.e., AvgSpeed map, is a 2D hexagonal binning plot quantifying average speed on the maze. 25 bins were used in x-direction and y-direction. To quantify similarity of the occupancy map and the average speed map between the training set and the test set, the 2D correlation coefficient (*r*) was used. If *r* = 1, it suggests that the training set and the test set have the same maps. A lower *r* indicates larger differences between two sets.

#### Baseline Decoder

For the baseline decoder, we used a feedforward neural network with a regression head, and identified ROIs as putative neurons from the microendoscopic images, and built a decoder based on temporal activities from these ROIs. Average fluorescence traces from N identified ROIs formed a N-dimensional input, and the decoder output the animal’s positions and running speed. Given that decoding performance depended on input representations, we prepared different representations as the following.

##### Smoothed traces of ROIs after background removal (FFNN(ROI))

We firfst obtained an average intensity projection from the image stacks, and then blurred the projection image using a moving 2D Gaussian filter with size of about 25% of the FOV, using ImageJ (Schindelin et al.,2012). The blurred projection was considered as the global background and is subtracted from all the raw images. Subsequently, an anisotropic diffusion operation (Perona and Malik, 1990) was applied to further denoise the images, followed by a morphological opening operation using a circular structure element with size similar to a neuron in the FOV. Afterwards, we obtained a maximum intensity projection from the denoised images whose backgrounds are removed, and annotated ROIs using the ImageJ Cell Magic Wand tool. The average fluorescence trace from each ROI is smoothed with a Savitzky-Golay smoothing filter.

##### Globally normalized denoised traces from CNMFe (FFNN(normG))

Another input representation was created by denoised temporal traces through applying the CNMFe algorithm (Zhou et al.,2018). The average fluorescence trace C from each ROI was normalized by the global maximum across all the traces. With this procedure, the correlations between the neurons were informative and thus relative magnitudes between them were preserved.

#### End-to-end decoder with image inputs

##### Regression task

The end-to-end decoder took each image frame as an input and predicted the animal’s behavioral attributes (positions and/or speed) at the corresponding time point. The decoding problem could be formulated as a multi-class classification task by binning the behavioral variables, but we chose to formulate the problem as a regression task for the following reasons. First, the animals were freely exploring the maze, so the maze occupancy and running speed distribution were highly imbalanced. However, the data size was not large enough for undersampling data to create a uniform distribution, yet oversampling minority classes using techniques like SMOTE (Chawla et al., 2002) introduced extra bias. In addition, the common loss function in a classification task could not keep the ordinal relationship between positions and speeds, i.e., prediction error between neighboring positions and error between distant positions were same using cross entropy.

The output dimensions (or number of attributes) depended on the decoding strategy (simultaneous or separate).

##### Simultaneous decoding

For simultaneous decoding strategy, convolutions and feed forward layers were shared to extract features for position and speed encoding, and different weights were used only in the final layer to predict position and speed. In this case, there are 3 output labels (*x, y, v*) in the regression head.

##### Separate decoding

For separate decoding strategy, there were two sets of convolutions and feed forward layers separately trained to predict position and speed. In this case, one decoder has 1 output label (*v*), and the other has 2 output labels (*x, y*).

##### Vanilla convolutional neural network (ConvNet)

The model took a grayscale (single channel) image as input and used convolutional layers connected to a regression head to decode continuous behaviour. A kernel size of 3 was used for all the 2D convolutions. The number of filters was doubled for each subsequent layer and saturated at 512. For downsampling, we used a stride of 2 to replace pooling layers for a better computational efficiency. After convolutions, features were flattened and fed into a series of dense layers. The number of units was halved for each subsequent dense layer. To encourage network sparsity and accelerate learning without a risk of introducing dying units, Leaky-Relu was used as the activation function except for the final layer. The final layer was applied with a Tanh activation function to ensure the output values match the range of target labels.

The number of convolutional layers, the number of fully connected layers, and the number of units in the first fully connected layer are hyperparameters. Evaluated on the validation set, the selected model used in this study had 8 convolutional layers and 5 fully connected layers (with number of units = [256, 64, 32, 16, 3] respectively). In our task, adding batch normalization made the prediction performance worse, so we did not use batch normalization in our decoder.

##### Residual neural network with 50 layers (ResNet-50)

The ResNet model (He et al., 2016) was originally designed to take a RGB image input for object recognition tasks, so the grayscale images were converted into RGB images, with the per-pixel mean subtracted. We kept the feature extraction architecture that was composed of convolutional blocks and identity blocks, and replaced the classification architecture with the dense layers whose activation functions were Leaky-Relu (the first few layers) and Tanh (the last layer).

For transfer learning, the parameters in feature extraction architecture were pre-trained by ImageNet and frozen during the model training, whereas the parameters in the dense layers are updated throughout training. Afterwards, the last few layers of the feature extraction architecture were unfrozen and jointly trained with the new dense layers. For learning from scratch, all the parameters in the model were initialized with Xavier initialization (Glorot and Bengio, 2010).

##### Efficient convolutional network for mobile applications (MobileNet)

MobileNet (Howard et al., 2017) was developed to achieve a lightweight architecture by replacing a convolutional layer with a depthwise convolutional layer, followed by a pointwise convolutional layer that combined these filtered values to create new features. It used a stride of 2 to reduce the spatial dimensions instead of pooling layers. Same preprocessing steps used in the ResNet model were applied to this decoder. Similarly, both the transfer learning and learning from scratch procedures were performed in this study.

##### Feedforward neural network (FFNN)

We built a feedforward neural network to learn features directly through dense connections. The model first applied a 2D max-pooling on each grayscale image for spatial downsampling, and then flattened these features for a series of dense layers. The number of units, activation functions, and procedure for hyperparameter searching were the same as in the vanilla ConvNet.

#### End-to-end decoder with video inputs

We also created an end-to-end decoder that took a short video with a window size of *N* to predict behavior at the corresponding time centroids. Two architectures were developed, depending on whether convolutional weights were shared across time.

##### Convolutions independent of time

The input dimension expanded from a single channel to multiple channels, each corresponding to an image frame. The decoder architecture kept the original vanilla convolutional neural network with 2D kernels. Features from different frames were weighted and integrated after the first layer. There was no constraint on convolutional kernels.

##### Convolutions shared across time

The input dimension expanded with a temporal dimension for *N* frames. Each frame went through a block of convolutional layers whose weights were shared across time, generating a frame-specific feature. These frame-specific features were linearly weighted by an attention layer, and then integrated into a final feature *F_post_* before feeding into the dense layers of the model.

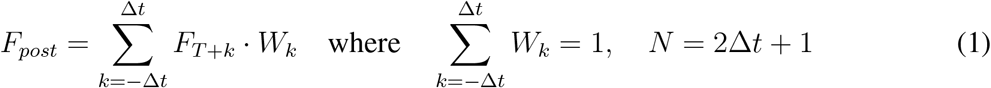

#### Training procedure

Mini-batch gradient descent was used for training the model. For each mini-batch, a subset of input images and output labels were randomly sampled from the training set. Each input image had corresponding normalized output labels (*x, y, v*). The batch size (i.e., number of samples per step) was 64 and the order of samples were further shuffled for each epoch. Hyperparameter searching was achieved by grid search for learning rate, number of the convolutional layers, number of the dense layers, number of units, and dropout. Adam optimization was used to accelerate learning. Kernel weights and biases are initialized by Xavier initialization (Glorot and Bengio, 2010), and updated by backpropagating the L1 loss, i.e., absolute differences between estimated and target values.

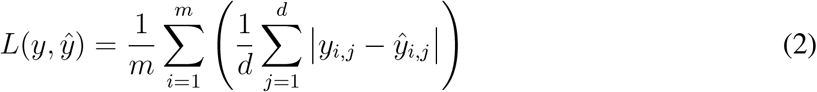

where *y_i_* is the ground truth, 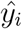 is the predicted value for sample *i, m* is the batch size, and *d* is the dimension of the model output (*d* =3 for simultaneous decoding and *d* =1 for separate decoding).

The training was stopped when the number of iteration steps reached 50,000 or when the validation performance remained unchanged or increased over 5 epochs. The model variables and training hyperparameters were saved in a checkpoint every 500 iteration steps. The training was performed on GeForce RTX 2080 Ti using Tensorflow, and parallel computation was utilized for grid search.

#### Evaluation metrics

After a model was trained, its performance was evaluated on the test set. Given that the model output normalized variables, the predicted values were transformed back to the original scales in [cm] and [cm/s]. Afterwards, we calculated the Euclidean distance between each ground truth and its predicted position, and used the median among these errors as a performance metric for position decoding. For speed decoding, we calculated the absolute difference between each ground truth and its predicted value, and used the median error as a performance metric. For an experiment or a specific model evaluated on several datasets, the distribution of the median decoding error was described as mean ± s.e.

##### Chance level calculation

The chance level was defined as decoding performance using Random images (see previous section Experiment on residual information).

### 5.5 Manifold analysis

To visualize the ensemble representation at each network layer in our decoder, ensembles at different time points were projected to a low-dimensional subspace.

#### Isomap

Isomap is a nonlinear dimensionality reduction algorithm for seeking embeddings that maintains geodesic distances between all data points. We used scikit-learn (Pedregosa et al., 2011) to implement this algorithm and extracted the low-dimensional embeddings of each layer. The optimal number of nearest neighbors (*K*_opt_) was selected by finding the elbow at which the reconstruction error (ER) curve ceased to decrease significantly with more nearest neighbors.

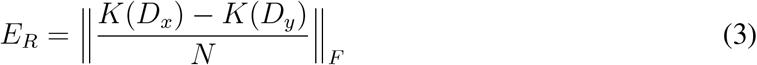

where *N* is the number of samples, *D_x_* and *D_y_* follow the same definitions above, and *K* is the Isomap kernel.

#### Manifold similarity metrics

To measure geometric similarity between the manifold and the behavioral topology formed by position, we calculated the Pearson correlation between pairwise distances in the 2D manifold coordinates and behavioral coordinates (Low et al., 2018), which was used as a metric to evaluate learning across different network layers.

### 5.6 Network saliency map

To better interpret how the encoder network makes the predictions, we used a saliency map to indicate the pixels whose changes have the most impact on the prediction. Instead of directly propagating the output gradient to the input (Simonyan et al., 2013), we modified the gradient-weighted class activation mapping (Grad-CAM) method (Selvaraju et al., 2017) to create a smoother saliency map.

Grad-CAM is originally designed to backpropagate the gradients of target class score *y^c^* to the last convolutional layer whose feature maps preserve spatial information, and generate a localized saliency map *L^c^* for class c, by a weighted combination of activation maps *A^k^* followed by ReLU operation, as in Eq (4).

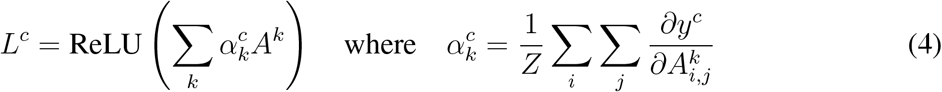

We modified Grad-CAM because our study used a continuous regression task. We visualized salience over input that either increased or decreased the output by computing the absolute gradients. We also adapted this algorithm to visualize saliency maps of different filters in the network model, by computing gradients from the filter at a specific layer with respect to the closest convolutional layer whose feature maps had large enough spatial resolution.

### 5.7 Statistical analysis

Statistical analyses were performed using SciPy (Virtanen et al., 2020). All values were reported as mean ± s.e. unless otherwise indicated. Wilcoxon signed-rank tests were used when paired samples were not normally distributed. For cases matching the normality requirement, Student’s t-test was used. Bonferroni correction was used for multiple comparisons. Pearson correlation was used to obtain correlations.

## References

J. Adebayo, J. Gilmer, M. Muelly, I. Goodfellow, M. Hardt, and B. Kim. Sanity checks for saliency maps. Advances in neural information processing systems, 31:9505–9515, 2018. 16

O. J. Ahmed and M. R. Mehta. Running speed alters the frequency of hippocampal gamma oscillations. Journal of Neuroscience, 32(21):7373–7383, 2012. 14

W. E. Allen, I. V. Kauvar, M. Z. Chen, E. B. Richman, S. J. Yang, K. Chan, V. Gradinaru, B. E. Deverman, L. Luo, and K. Deisseroth. Global representations of goal-directed behavior in distinct cell types of mouse neocortex. Neuron, 94(4):891–907, 2017. 16

E. W. Archer, U. Koster, J. W. Pillow, and J. H. Macke. Low-dimensional models of neural population activity in sensory cortical circuits. Advances in neural information processing systems, 27:343–351, 2014. 11

B. B. Averbeck, P. E. Latham, and A. Pouget. Neural correlations, population coding and computation. Nature reviews neuroscience, 7(5):358–366, 2006. 4

J. N. Betley, S. Xu, Z. F. H. Cao, R. Gong, C. J. Magnus, Y. Yu, and S. M. Sternson. Neurons for hunger and thirst transmit a negative-valence teaching signal. Nature, 521(7551):180–185, 2015. 1

K. C. Bittner, C. Grienberger, S. P. Vaidya, A. D. Milstein, J. J. Macklin, J. Suh, S. Tonegawa, and J. C. Magee. Conjunctive input processing drives feature selectivity in hippocampal ca1 neurons. Nature neuroscience, 18(8):1133–1142, 2015. 16

G. J. Broussard, Y. Liang, M. Fridman, E. K. Unger, G. Meng, X. Xiao, N. Ji, L. Petreanu, and L. Tian. In vivo measurement of afferent activity with axon-specific calcium imaging. Nature neuroscience, 21(9):1272–1280, 2018. 16

E. N. Brown, L. M. Frank, D. Tang, M. C. Quirk, and M. A. Wilson. A statistical paradigm for neural spike train decoding applied to position prediction from ensemble firing patterns of rat hippocampal place cells. Journal of Neuroscience, 18(18):7411–7425, 1998. 3, 16

D. J. Cai, D. Aharoni, T. Shuman, J. Shobe, J. Biane, W. Song, B. Wei, M. Veshkini, M. La-Vu, J. Lou, et al. A shared neural ensemble links distinct contextual memories encoded close in time. Nature, 534(7605):115–118, 2016. 2

L. Cao, V. Varga, and Z. S. Chen. Ultrafast readout of representations from spatially distributed rodent hippocampal field potentials. bioRxiv, page 828467, 2019. 16

J. M. Carmena, M. A. Lebedev, R. E. Crist, J. E. O’Doherty, D. M. Santucci, D. F. Dimitrov, P. G. Patil, C. S. Henriquez, and M. A. Nicolelis. Learning to control a brain–machine interface for reaching and grasping by primates. PLoS biol, 1(2):e42, 2003. 3

N. V. Chawla, K. W. Bowyer, L. O. Hall, and W. P. Kegelmeyer. Smote: synthetic minority over-sampling technique. Journal of artificial intelligence research, 16:321–357, 2002. 38

M. M. Churchland, J. P. Cunningham, M. T. Kaufman, J. D. Foster, P. Nuyujukian, S. I. Ryu, and K. V. Shenoy. Neural population dynamics during reaching. Nature, 487(7405):51–56, 2012. 11

C. J. Cueva, A. Saez, E. Marcos, A. Genovesio, M. Jazayeri, R. Romo, C. D. Salzman, M. N. Shadlen, and S. Fusi. Low-dimensional dynamics for working memory and time encoding. Proceedings of the National Academy of Sciences, 117(37):23021–23032, 2020. 11

T. J. Davidson, F. Kloosterman, and M. A. Wilson. Hippocampal replay of extended experience. Neuron, 63(4):497–507, 2009. 14

X. Deng, D. F. Liu, K. Kay, L. M. Frank, and U. T. Eden. Clusterless decoding of position from multiunit activity using a marked point process filter. Neural computation, 27(7):1438–1460, 2015. 16

A. Dubbs, J. Guevara, and R. Yuste. moco: Fast motion correction for calcium imaging. Frontiers in neuroinformatics, 10:6, 2016. 3

T. Ebina, Y. Masamizu, Y. R. Tanaka, A. Watakabe, R. Hirakawa, Y. Hirayama, R. Hira, S.-I. Terada, D. Koketsu, K. Hikosaka, et al. Two-photon imaging of neuronal activity in motor cortex of marmosets during upper-limb movement tasks. Nature communications, 9(1):1–16, 2018. 1

G. Etter, F. Manseau, and S. Williams. A probabilistic framework for decoding behavior from in vivo calcium imaging data. Frontiers in Neural Circuits, 14:19, 2020. 16

T. Euler, P. B. Detwiler, and W. Denk. Directionally selective calcium signals in dendrites of starburst amacrine cells. Nature, 418(6900):845–852, 2002. 15

D. A. Evans, A. V. Stempel, R. Vale, S. Ruehle, Y. Lefler, and T. Branco. A synaptic threshold mechanism for computing escape decisions. Nature, 558(7711):590–594, 2018. 1

B. A. Flusberg, A. Nimmerjahn, E. D. Cocker, E. A. Mukamel, R. P. Barretto, T. H. Ko, L. D. Burns, J. C. Jung, and M. J. Schnitzer. High-speed, miniaturized fluorescence microscopy in freely moving mice. Nature methods, 5(11):935–938, 2008. 2

M. Frey, S. Tanni, C. Perrodin, A. O’Leary, M. Nau, J. Kelly, A. Banino, C. F. Doeller, and C. Barry. Deepinsight: a general framework for interpreting wide-band neural activity. bioRxiv, page 871848, 2019. 4, 16

E. Ganmor, M. Krumin, L. F. Rossi, M. Carandini, and E. P. Simoncelli. Direct estimation of firing rates from calcium imaging data. arXiv preprint arXiv:1601.00364, 2016. 16

C. Geisler, D. Robbe, M. Zugaro, A. Sirota, and G. Buzsáki. Hippocampal place cell assemblies are speed-controlled oscillators. Proceedings of the National Academy of Sciences, 104(19): 8149–8154, 2007. 14

A. P. Georgopoulos, A. B. Schwartz, and R. E. Kettner. Neuronal population coding of movement direction. Science, 233(4771):1416–1419, 1986. 3

K. K. Ghosh, L. D. Burns, E. D. Cocker, A. Nimmerjahn, Y. Ziv, A. El Gamal, and M. J. Schnitzer. Miniaturized integration of a fluorescence microscope. Nature methods, 8(10):871, 2011. 2

A. Glas, M. Hübener, T. Bonhoeffer, and P. M. Goltstein. Benchmarking miniaturized microscopy against two-photon calcium imaging using single-cell orientation tuning in mouse visual cortex. PloS one, 14(4):e0214954, 2019. 1

J. I. Glaser, A. S. Benjamin, R. H. Chowdhury, M. G. Perich, L. E. Miller, and K. P. Kording. Machine learning for neural decoding. Eneuro, 7(4), 2020. 3, 16

X. Glorot and Y. Bengio. Understanding the difficulty of training deep feedforward neural networks. In Proceedings of the thirteenth international conference on artificial intelligence and statistics, pages 249–256, 2010. 39, 40

W. Gobel and F. Helmchen. In vivo calcium imaging of neural network function. Physiology, 22(6): 358–365, 2007. 15

Z. H. T. Góis and A. B. Tort. Characterizing speed cells in the rat hippocampus. Cell reports, 25(7): 1872–1884, 2018. 13

W. G. Gonzalez, H. Zhang, A. Harutyunyan, and C. Lois. Persistence of neuronal representations through time and damage in the hippocampus. Science, 365(6455):821–825, 2019. 16

B. F. Grewe, J. Gründemann, L. J. Kitch, J. A. Lecoq, J. G. Parker, J. D. Marshall, M. C. Larkin, P. E. Jercog, F. Grenier, J. Z. Li, et al. Neural ensemble dynamics underlying a long-term associative memory. Nature, 543(7647):670–675, 2017. 1

C. D. Harvey, P. Coen, and D. W. Tank. Choice-specific sequences in parietal cortex during a virtual-navigation decision task. Nature, 484(7392):62–68, 2012. 1

K. He, X. Zhang, S. Ren, and J. Sun. Deep residual learning for image recognition. In Proceedings of the IEEE conference on computer vision and pattern recognition, pages 770–778, 2016. 39

F. Helmchen, M. S. Fee, D. W. Tank, and W. Denk. A miniature head-mounted two-photon microscope: high-resolution brain imaging in freely moving animals. Neuron, 31(6):903–912, 2001. 2

A. G. Howard, M. Zhu, B. Chen, D. Kalenichenko, W. Wang, T. Weyand, M. Andreetto, and H. Adam. Mobilenets: Efficient convolutional neural networks for mobile vision applications. arXiv preprint arXiv:1704.04861, 2017. 39

D. Huber, D. A. Gutnisky, S. Peron, D. H. O’connor, J. S. Wiegert, L. Tian, T. G. Oertner, L. L. Looger, and K. Svoboda. Multiple dynamic representations in the motor cortex during sensorimotor learning. Nature, 484(7395):473–478, 2012. 1

J. H. Jennings, R. L. Ung, S. L. Resendez, A. M. Stamatakis, J. G. Taylor, J. Huang, K. Veleta, P. A. Kantak, M. Aita, K. Shilling-Scrivo, et al. Visualizing hypothalamic network dynamics for appetitive and consummatory behaviors. Cell, 160(3):516–527, 2015. 1

S. Jewell and D. Witten. Exact spike train inference via l0 optimization. The annals of applied statistics, 12(4):2457, 2018. 3

P. Kaifosh, J. D. Zaremba, N. B. Danielson, and A. Losonczy. Sima: Python software for analysis of dynamic fluorescence imaging data. Frontiers in neuroinformatics, 8:80, 2014. 3

S. W. Keemink, S. C. Lowe, J. M. Pakan, E. Dylda, M. C. Van Rossum, and N. L. Rochefort. Fissa: A neuropil decontamination toolbox for calcium imaging signals. Scientific reports, 8(1):1–12, 2018. 15

J. N. Kerr, D. Greenberg, and F. Helmchen. Imaging input and output of neocortical networks in vivo. Proceedings of the National Academy of Sciences, 102(39):14063–14068, 2005. 15

A. Klaus, G. J. Martins, V. B. Paixao, P. Zhou, L. Paninski, and R. M. Costa. The spatiotemporal organization of the striatum encodes action space. Neuron, 95(5):1171–1180, 2017. 1, 3

T. Kleindienst, J. Winnubst, C. Roth-Alpermann, T. Bonhoeffer, and C. Lohmann. Activitydependent clustering of functional synaptic inputs on developing hippocampal dendrites. Neuron, 72(6):1012–1024, 2011. 15

F. Kloosterman, S. P. Layton, Z. Chen, and M. A. Wilson. Bayesian decoding using unsorted spikes in the rat hippocampus. Journal of neurophysiology, 2014. 16

E. Kropff, J. E. Carmichael, M.-B. Moser, and E. I. Moser. Speed cells in the medial entorhinal cortex. Nature, 523(7561):419–424, 2015. 13

M. E. Larkum, K. Kaiser, and B. Sakmann. Calcium electrogenesis in distal apical dendrites of layer 5 pyramidal cells at a critical frequency of back-propagating action potentials. Proceedings of the National Academy of Sciences, 96(25):14600–14604, 1999. 15

M. Lavzin, S. Rapoport, A. Polsky, L. Garion, and J. Schiller. Nonlinear dendritic processing determines angular tuning of barrel cortex neurons in vivo. Nature, 490(7420):397–401, 2012. 15

S. Lee, J. F. Meyer, J. Park, and S. M. Smirnakis. Visually driven neuropil activity and information encoding in mouse primary visual cortex. Frontiers in neural circuits, 11:50, 2017. 16

G. Lopes, N. Bonacchi, J. Frazão, J. P. Neto, B. V. Atallah, S. Soares, L. Moreira, S. Matias, P. M. Itskov, P. A. Correia, et al. Bonsai: an event-based framework for processing and controlling data streams. Frontiers in neuroinformatics, 9:7, 2015. 35

R. J. Low, S. Lewallen, D. Aronov, R. Nevers, and D. W. Tank. Probing variability in a cognitive map using manifold inference from neural dynamics. bioRxiv, page 418939, 2018. 41

J. Lu, C. Li, J. Singh-Alvarado, Z. C. Zhou, F. Fröhlich, R. Mooney, and F. Wang. Min1pipe: A miniscope 1-photon-based calcium imaging signal extraction pipeline. Cell reports, 23(12): 3673–3684, 2018. 2, 3, 4, 15

V. Mante, D. Sussillo, K. V. Shenoy, and W. T. Newsome. Context-dependent computation by recurrent dynamics in prefrontal cortex. nature, 503(7474):78–84, 2013. 11

W. Mittmann, D. J. Wallace, U. Czubayko, J. T. Herb, A. T. Schaefer, L. L. Looger, W. Denk, and J. N. Kerr. Two-photon calcium imaging of evoked activity from l5 somatosensory neurons in vivo. Nature neuroscience, 14(8):1089–1093, 2011. 1

E. A. Mukamel, A. Nimmerjahn, and M. J. Schnitzer. Automated analysis of cellular signals from large-scale calcium imaging data. Neuron, 63(6):747–760, 2009. 3

H. Nover, C. H. Anderson, and G. C. DeAngelis. A logarithmic, scale-invariant representation of speed in macaque middle temporal area accounts for speed discrimination performance. Journal of Neuroscience, 25(43):10049–10060, 2005. 13

K. Ohki, S. Chung, Y. H. Ch’ng, P. Kara, and R. C. Reid. Functional imaging with cellular resolution reveals precise micro-architecture in visual cortex. Nature, 433(7026):597–603, 2005. 15

H. F. Ólafsdóttir, F. Carpenter, and C. Barry. Task demands predict a dynamic switch in the content of awake hippocampal replay. Neuron, 96(4):925–935, 2017. 14

M. Pachitariu, C. Stringer, M. Dipoppa, S. Schröder, L. F. Rossi, H. Dalgleish, M. Carandini, and K. D. Harris. Suite2p: beyond 10,000 neurons with standard two-photon microscopy. Biorxiv, 2017. 15

F. Pedregosa, G. Varoquaux, A. Gramfort, V. Michel, B. Thirion, O. Grisel, M. Blondel, P. Pretten-hofer, R. Weiss, V. Dubourg, J. Vanderplas, A. Passos, D. Cournapeau, M. Brucher, M. Perrot, and E. Duchesnay. Scikit-learn: Machine learning in Python. Journal of Machine Learning Research, 12:2825–2830, 2011. 41

F. Pereira, T. Mitchell, and M. Botvinick. Machine learning classifiers and fmri: a tutorial overview. Neuroimage, 45(1):S199–S209, 2009. 3

P. Perona and J. Malik. Scale-space and edge detection using anisotropic diffusion. IEEE Transactions on pattern analysis and machine intelligence, 12(7):629–639, 1990. 38

L. Petreanu, D. A. Gutnisky, D. Huber, N.-l. Xu, D. H. O’Connor, L. Tian, L. Looger, and K. Svoboda. Activity in motor–sensory projections reveals distributed coding in somatosensation. Nature, 489(7415):299–303, 2012. 16

L. Pinto and Y. Dan. Cell-type-specific activity in prefrontal cortex during goal-directed behavior. Neuron, 87(2):437–450, 2015. 1, 3

E. A. Pnevmatikakis. Analysis pipelines for calcium imaging data. Current opinion in neurobiology, 55:15–21, 2019. 2

E. A. Pnevmatikakis and A. Giovannucci. Normcorre: An online algorithm for piecewise rigid motion correction of calcium imaging data. Journal of neuroscience methods, 291:83–94, 2017. 3, 35

E. A. Pnevmatikakis, D. Soudry, Y. Gao, T. A. Machado, J. Merel, D. Pfau, T. Reardon, Y. Mu, C. Lacefield, W. Yang, et al. Simultaneous denoising, deconvolution, and demixing of calcium imaging data. Neuron, 89(2):285–299, 2016. 3, 15

R. Q. Quiroga and S. Panzeri. Extracting information from neuronal populations: information theory and decoding approaches. Nature Reviews Neuroscience, 10(3):173–185, 2009. 3

J. Reidl, J. Starke, D. B. Omer, A. Grinvald, and H. Spors. Independent component analysis of high-resolution imaging data identifies distinct functional domains. Neuroimage, 34(1):94–108, 2007. 3

T. F. Roberts, E. Hisey, M. Tanaka, M. G. Kearney, G. Chattree, C. F. Yang, N. M. Shah, and R. Mooney. Identification of a motor-to-auditory pathway important for vocal learning. Nature neuroscience, 20(7):978, 2017. 1

G. Rothschild, I. Nelken, and A. Mizrahi. Functional organization and population dynamics in the mouse primary auditory cortex. Nature neuroscience, 13(3):353, 2010. 1

E. Salinas and L. Abbott. Vector reconstruction from firing rates. Journal of computational neuroscience, 1(1-2):89–107, 1994. 3

T. D. Sanger. Probability density estimation for the interpretation of neural population codes. Journal of neurophysiology, 76(4):2790–2793, 1996. 3

J. Sawinski, D. J. Wallace, D. S. Greenberg, S. Grossmann, W. Denk, and J. N. Kerr. Visually evoked activity in cortical cells imaged in freely moving animals. Proceedings of the National Academy of Sciences, 106(46):19557–19562, 2009. 2

J. Schindelin, I. Arganda-Carreras, E. Frise, V. Kaynig, M. Longair, T. Pietzsch, S. Preibisch, C. Rueden, S. Saalfeld, B. Schmid, et al. Fiji: an open-source platform for biological-image analysis. Nature methods, 9(7):676–682, 2012. 38

R. R. Selvaraju, M. Cogswell, A. Das, R. Vedantam, D. Parikh, and D. Batra. Grad-cam: Visual explanations from deep networks via gradient-based localization. In Proceedings of the IEEE international conference on computer vision, pages 618–626, 2017. 10, 16, 41

M. E. Sheffield and D. A. Dombeck. Calcium transient prevalence across the dendritic arbour predicts place field properties. Nature, 517(7533):200–204, 2015. 16

O. A. Shemesh, C. Linghu, K. D. Piatkevich, D. Goodwin, O. T. Celiker, H. J. Gritton, M. F. Romano, R. Gao, C.-C. J. Yu, H.-A. Tseng, et al. Precision calcium imaging of dense neural populations via a cell-body-targeted calcium indicator. Neuron, 107(3):470–486, 2020. 16

T. Shuman, D. Aharoni, D. J. Cai, C. R. Lee, S. Chavlis, L. Page-Harley, L. M. Vetere, Y. Feng, C. Y. Yang, I. Mollinedo-Gajate, et al. Breakdown of spatial coding and interneuron synchronization in epileptic mice. Nature neuroscience, 23(2):229–238, 2020. 16

K. Simonyan, A. Vedaldi, and A. Zisserman. Deep inside convolutional networks: Visualising image classification models and saliency maps. arXiv preprint arXiv:1312.6034, 2013. 16, 41

B. Sivyer and S. R. Williams. Direction selectivity is computed by active dendritic integration in retinal ganglion cells. Nature neuroscience, 16(12):1848–1856, 2013. 15

S. L. Smith, I. T. Smith, T. Branco, and M. Häusser. Dendritic spikes enhance stimulus selectivity in cortical neurons in vivo. Nature, 503(7474):115–120, 2013. 15

J. T. Springenberg, A. Dosovitskiy, T. Brox, and M. Riedmiller. Striving for simplicity: The all convolutional net. arXiv preprint arXiv:1412.6806, 2014. 16

F. Stefanini, L. Kushnir, J. C. Jimenez, J. H. Jennings, N. I. Woods, G. D. Stuber, M. A. Kheirbek, R. Hen, and S. Fusi. A distributed neural code in the dentate gyrus and in ca1. Neuron, 2020. 16

K. Svoboda, F. Helmchen, W. Denk, and D. W. Tank. Spread of dendritic excitation in layer 2/3 pyramidal neurons in rat barrel cortex in vivo. Nature neuroscience, 2(1):65–73, 1999. 15

N. Takahashi, K. Kitamura, N. Matsuo, M. Mayford, M. Kano, N. Matsuki, and Y. Ikegaya. Locally synchronized synaptic inputs. Science, 335(6066):353–356, 2012. 15

A. Tampuu, T. Matiisen, H. F. Ólafsdóttir, C. Barry, and R. Vicente. Efficient neural decoding of self-location with a deep recurrent network. PLoS computational biology, 15(2):e1006822, 2019. 16

Y. Tanimoto, A. Yamazoe-Umemoto, K. Fujita, Y. Kawazoe, Y. Miyanishi, S. J. Yamazaki, X. Fei, K. E. Busch, K. Gengyo-Ando, J. Nakai, et al. Calcium dynamics regulating the timing of decision-making in c. elegans. Elife, 6:e21629, 2017. 1

M. Tu, R. Zhao, A. Adler, W.-B. Gan, and Z. S. Chen. Efficient position decoding methods based on fluorescence calcium imaging in the mouse hippocampus. Neural Computation, 32(6):1144–1167, 2020. 16

P. Virtanen, R. Gommers, T. E. Oliphant, M. Haberland, T. Reddy, D. Cournapeau, E. Burovski, P. Peterson, W. Weckesser, J. Bright, S. J. van der Walt, M. Brett, J. Wilson, K. J. Millman, N. Mayorov, A. R. J. Nelson, E. Jones, R. Kern, E. Larson, C. J. Carey, İ. Polat, Y. Feng, E. W. Moore, J. VanderPlas, D. Laxalde, J. Perktold, R. Cimrman, I. Henriksen, E. A. Quintero, C. R. Harris, A. M. Archibald, A. H. Ribeiro, F. Pedregosa, P. van Mulbregt, and SciPy 1.0 Contributors. SciPy 1.0: Fundamental Algorithms for Scientific Computing in Python. Nature Methods, 17: 261–272, 2020. doi: 10.1038/s41592-019-0686-2. 41

J. T. Vogelstein, A. M. Packer, T. A. Machado, T. Sippy, B. Babadi, R. Yuste, and L. Paninski. Fast nonnegative deconvolution for spike train inference from population calcium imaging. Journal of neurophysiology, 104(6):3691–3704, 2010. 3

M. Wang, X. Liao, R. Li, S. Liang, R. Ding, J. Li, J. Zhang, W. He, K. Liu, J. Pan, et al. Singleneuron representation of learned complex sounds in the auditory cortex. Nature communications, 11(1):1–14, 2020. 1

Z. Wang, A. C. Bovik, H. R. Sheikh, and E. P. Simoncelli. Image quality assessment: from error visibility to structural similarity. IEEE transactions on image processing, 13(4):600–612, 2004. 37

M. A. Wilson and B. L. McNaughton. Dynamics of the hippocampal ensemble code for space. Science, 261(5124):1055–1058, 1993. 3, 16

W. Wu, Y. Gao, E. Bienenstock, J. P. Donoghue, and M. J. Black. Bayesian population decoding of motor cortical activity using a kalman filter. Neural computation, 18(1):80–118, 2006. 3

E. Yaksi and R. W. Friedrich. Reconstruction of firing rate changes across neuronal populations by temporally deconvolved ca 2+ imaging. Nature methods, 3(5):377–383, 2006. 3

T. Yoshida and K. Ohki. Natural images are reliably represented by sparse and variable populations of neurons in visual cortex. Nature communications, 11(1):1–19, 2020. 1

K. Yu, S. Ahrens, X. Zhang, H. Schiff, C. Ramakrishnan, L. Fenno, K. Deisseroth, F. Zhao, M.-H. Luo, L. Gong, et al. The central amygdala controls learning in the lateral amygdala. Nature neuroscience, 20(12):1680–1685, 2017. 1

R. Yuste and W. Denk. Dendritic spines as basic functional units of neuronal integration. Nature, 375(6533):682–684, 1995. 15

M. D. Zeiler and R. Fergus. Visualizing and understanding convolutional networks. In European conference on computer vision, pages 818–833. Springer, 2014. 16

K. Zhang, I. Ginzburg, B. L. McNaughton, and T. J. Sejnowski. Interpreting neuronal population activity by reconstruction: unified framework with application to hippocampal place cells. Journal of neurophysiology, 79(2):1017–1044, 1998. 3, 16

P. Zhou, S. L. Resendez, J. Rodriguez-Romaguera, J. C. Jimenez, S. Q. Neufeld, A. Giovannucci, J. Friedrich, E. A. Pnevmatikakis, G. D. Stuber, R. Hen, et al. Efficient and accurate extraction of in vivo calcium signals from microendoscopic video data. Elife, 7:e28728, 2018. 2, 3, 4, 15, 36, 38

Y. Ziv, L. D. Burns, E. D. Cocker, E. O. Hamel, K. K. Ghosh, L. J. Kitch, A. El Gamal, and M. J. Schnitzer. Long-term dynamics of ca1 hippocampal place codes. Nature neuroscience, 16(3):264, 2013. 1, 16

W. Zong, R. Wu, M. Li, Y. Hu, Y. Li, J. Li, H. Rong, H. Wu, Y. Xu, Y. Lu, et al. Fast high-resolution miniature two-photon microscopy for brain imaging in freely behaving mice. Nature methods, 14(7):713–719, 2017. 2

